# Spatially structured host genetic diversity leads to the evolution of local specialization

**DOI:** 10.1101/2023.05.12.540593

**Authors:** Elisa Visher, Anisha Ali, Jonathan Barajas, Sehar Masud, Annika McBride, Edwin Ramos, Melissa Sui, Cristina Villalobos-Heredia, Natalie Walzer, P. Signe White, Mike Boots

## Abstract

Host heterogeneity and spatial population structure each influence parasite evolution but may also interact because space structures contacts between host types. Here, we experimentally evolve granulosis virus in microcosms of its *Plodia interpunctella* (Indian meal moth) host that differ in both spatial structure and host genetic diversity. We control spatial structure by manipulating the viscosity of the food that the larvae live within and host genetic diversity by adding larvae from either a single or two inbred lines to opposite ends of the microcosm. We preserve spatial structure across passages and assay virus from different positions within the microcosm on both host genotypes. We find that the lower contact rates between host genotypes resulting from spatial structure can lead to the evolution of locally specialized virus, even when the host population is genetically diverse overall. We also find that spatial structure changes how viruses specialize: viruses evolved in well-mixed environments had lower exploitation of familiar hosts, while those in spatially structured environments exhibited higher exploitation of familiar hosts. These results demonstrate that spatial structure and host heterogeneity interact to shape pathogen specialization and that the evolutionary consequences of host diversity depends on the population structure.

## INTRODUCTION

A body of literature on the ‘monoculture effect’ has shown that host genetic diversity affects infectious disease dynamics in both ecological and evolutionary contexts [1–5]. Specifically, genetically diverse host populations can decrease the prevalence of infectious diseases and may constrain their evolution of transmissibility or virulence [3,6–12]. This can be explained by heterogeneous host populations limiting the transmission of specialist pathogens and thereby selecting for ‘mediocre’ generalists that may be constrained by costs associated with expanded niche breadth [13–19]. This theory, however, only considers scenarios where the genetic diversity of the host population is well-mixed so that transmission occurs readily between host types. However, it is clear that many host populations exhibit some sort of spatial or contact network structuring. In this case, a parasite lineage may be more likely to infect one host type even if the population is genetically diverse overall. The impact of this interaction between host genetic diversity and spatial population structure on the evolution of pathogens is not well understood.

Several theoretical studies have examined the impact of spatial structure by manipulating the contact matrix between hosts to show that the degree of mixing in a population should change the evolutionarily optimal strategy for host range [20–22]. Higher contact rates between host types select for generalists in simple models [20–22], but can theoretically also lead to greater specialization when migration increases the amount of genetic variation that selection can act upon or increases local competition for patches [23–25]. Additionally, spatially structured transmission may impose additional selection pressures beyond the effect on host type mixing. Theory and experiments show that more local transmission results in ‘self-shading’ that can select for less infective, more prudent pathogens due to both genetic and ecological correlations that arise in spatial populations [26–34], though intermediate spatial structure can also select for higher exploitation rates [28,33]. Finally, these host range trade-offs and life history trade-offs may interact. Theory considering multidimensional trade-offs has shown that interactions between trade-offs of different classes can produce evolutionary outcomes that deviate from the predictions of single trade-off models [35–37]. This means that how organisms evolve to specialize or may also depend on the selective pressures acting on other trade-offs [35].

There are, therefore, many open questions for how we expect spatial structure and host genetic diversity to interact, both in terms of how host type mixing affects host range evolution and also in terms of how these different selection pressures interact. We explore these interactions by experimentally evolving granulosis virus in microcosms of its *Plodia interpunctella*, or Indian meal moth (Hübner), host that vary in their host genetic diversity and spatial structuring. We alter host heterogeneity by adding larvae from either the same or different inbred populations (host genotypes) to either half of the microcosm [38] (Figure 1). We manipulate spatial structure by altering the viscosity of the food medium that larvae live within [30,39,40]. Food viscosity therefore both manipulates the contact rates between hosts and structures infectious contacts [20,30]. The treatments are then fully crossed so that each host treatment is in both food viscosity treatments. Our experiment therefore explores how spatial structure and host heterogeneity interact and how the spatial structuring of heterogeneity affects its impact on pathogen evolution. We demonstrate that spatially structured host heterogeneity can promote local specialization and constrain the evolution of generalist viruses. Moreover, altering the spatial structure of a population changes how specialization affects pathogen traits. Specifically, the direction of specialization’s effect on host exploitation rate—whether pathogens exhibit higher or lower exploitation on their familiar host compared to a foreign host—depends on the spatial structure under which they evolved.

**Figure 1:**
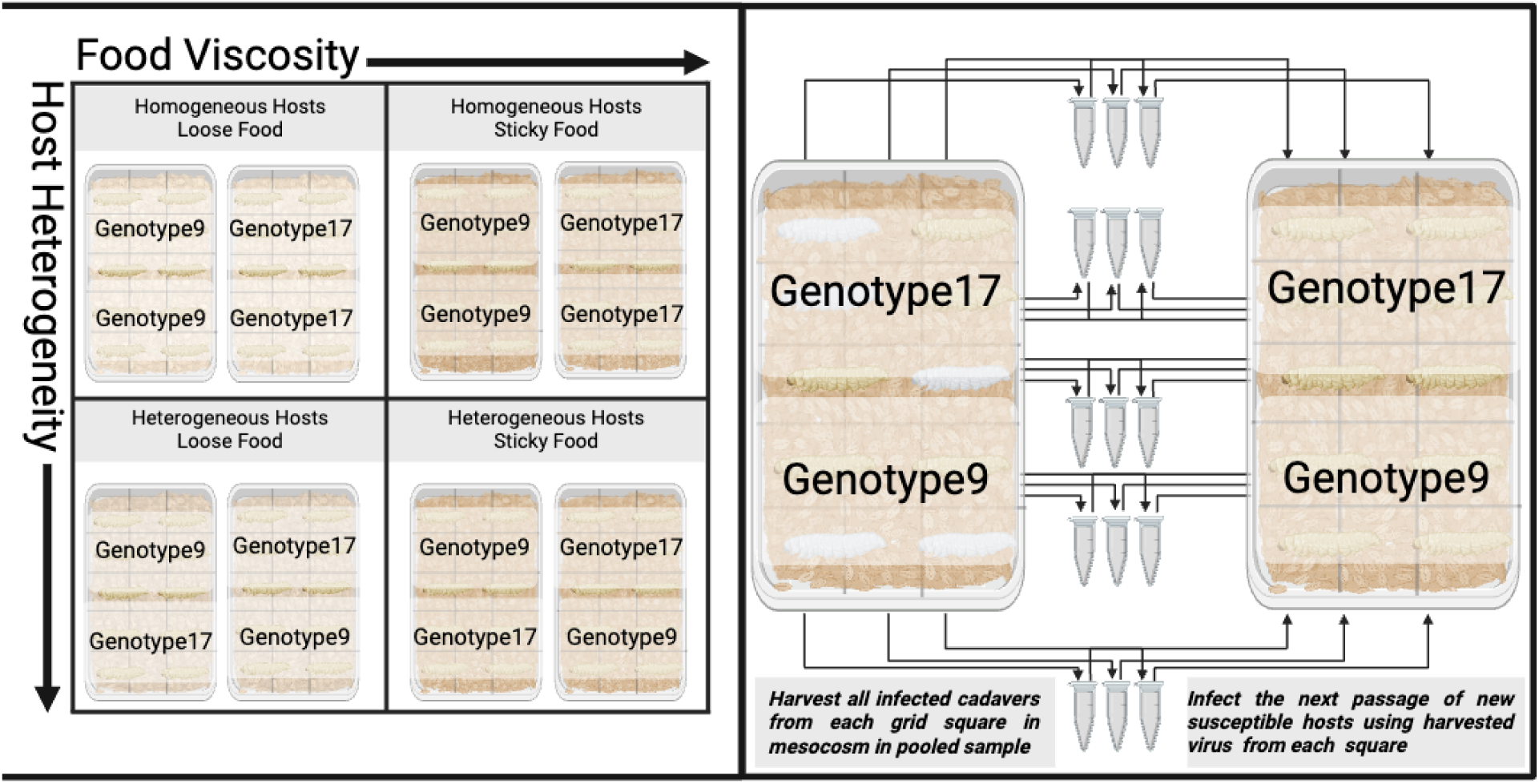
Experimental design and passaging scheme. (Left) Each microcosm was set up by adding 2 pots of food containing 11-day old larvae to opposite vertical ends of a plastic microcosm with a 3x5 grid. The 3^rd^ row of the grid is therefore a contact zone between the hosts. There are 3 replicates for each of the 8 microcosms. There are 4 different treatment types of homogeneous or heterogeneous hosts by ‘loose’ or ‘sticky’ food. Each homogeneous treatment was set up with both host types and each heterogeneous host treatment was set up alternating whic host was on ‘top’ or ‘bottom’ of the grid. Microcosms were consistently oriented in the growth chamber so that the ‘bottom’ side was closer to the door. (Right) After 20 days incubating with virus, microcosms were frozen and dissected into grid squares. All infected larvae from each grid square were harvested, added to a 1.5mL Eppendorf tube (1 square per tube), and homogenized to release occlusion bodies. 500mL of the virus solution was then pipetted in droplets over the same grid square of a new microcosm of the same treatment set-up with fresh 11-day old hosts. Therefore, the spatial force of infection was preserved across passages. We experimentally evolved virus for 10 passages.

## METHODS

### Study system

*Plodia interpunctella*, or the Indian meal moth, is a stored grain pest that naturally lives at high population density within its cereal food medium for 5 larval instar stages before it pupates into an adult moth that disperses, mates, and lays eggs, but does not eat [41,42]. It has a semelparous, monthly cyclical life cycle where the population moves through life stages roughly concordantly [40]. It is naturally infected by the *Plodia interpunctella* granulosis virus (PiGV), a dsDNA baculovirus that transmits when larvae directly cannibalize infection killed cadavers or consume virus that was released into the environment from such cadavers [43,44]. Thus, the virus is an obligate killer that will only transmit if it kills its host. Notably, the virus can only infect *P. interpunctella* during the larval stages and, if exposed larvae clear the infection, they can pupate and carry out the rest of their life history as normal [45]. Infection killed cadavers are recognizable because successful infection turns larvae an opaque, chalky white color due to the high density of viral occlusion bodies [46]. The system has proved a powerful tool for examining the effects of spatial structure on pathogen exploitation rates and, more recently, host genotype specialization [30,38].

### Host genotype selection and maintenance

For this experiment, we selected two inbred populations of *P. interpunctella* that were previously generated in the lab by brother-sister mating individuals for >27 generations [47]. Each population should therefore be essentially a clonal population of two distinct host genotypes (Genotype 9 and Genotype 17). These genotypes were chosen because they had similar levels of general resistance or, alternatively worded, the ancestral virus has similar levels of initial infectivity on them [38] (Table S2, Figure S3). We have previously shown that virus evolved in homogeneous populations of these specific host genotypes can specialize on its familiar host genotype by increasing their infectivity and/or productivity [38].

We maintained these genotypes as inbred lines in wide-mouthed Nalgene pots (ThermoFisher Scientific, U.K.) in the absence of infection with 200g ‘Standard’ food medium (made in batches of 500g cereal mix (50% Earth’s Best oatmeal, 30% wheat bran, 20% rice flour), 100g brewer’s yeast, 2.2g sorbic acid, 2.2g methyl paraben, 125ml honey, and 125ml glycerol) in incubators set at 27±2 °C and 35±5% humidity, with 16:8hr light: dark cycles [48]. To maintain populations, ∼50 adult moths were moved into a new pot with new food when they emerged monthly. Thus, host lines were maintained in the absence of infection and did not evolve throughout the experiment.

### Manipulating spatial structure

We manipulated spatial structure in the experiment by changing the viscosity of the food medium as in [30,39,40]. Because larvae live within their food medium, this alters individuals’ dispersal patterns and thus the spatial structure of their infectious contacts. We made food of two viscosities, ‘loose’ and ‘sticky’, which differ solely in the amount of glycerol added. ‘Loose’ food contains 500g Earth’s Best oatmeal, 100g brewer’s yeast, 2.2g sorbic acid, 2.2g methyl paraben, and 175ml glycerol. ‘Sticky’ food contains 500g Earth’s Best oatmeal, 100g brewer’s yeast, 2.2g sorbic acid, 2.2g methyl paraben, and 450ml glycerol. ‘Loose’ and ‘sticky’ food wa then added to wide-mouthed Nalgene pots and frozen overnight to kill any insect eggs potentially contaminating the food. The mass of food added to these pots (300g ‘loose’ and 550g ‘sticky’ food) was determined by the volume needed to entirely fill half of one of our plastic microcosm containers.

We tested the impact of food viscosity on spatial structure in several ways. First, we measured life history traits (development time and mass at pupation) of larvae added to assay grids with ‘sticky’ and ‘loose’ food (as in [48]) to ensure that food viscosity did not affect these metric (Figure S1). Second, we added ‘loose’ and ‘sticky’ food to 4 cm diameter PVC pipes (10 each), added 11-day old 3rd instar larvae to one side of the pipe, and measured how far larvae dispersed after 10 days (Table S1). Finally, we set up microcosms with the two food types as in our experiment below but added food containing larvae to only one half of the microcosm and sterile food to the other half. We incubated these microcosms for 21 days, then recorded the position of every individual (Figure 2A).

**Figure 2:**
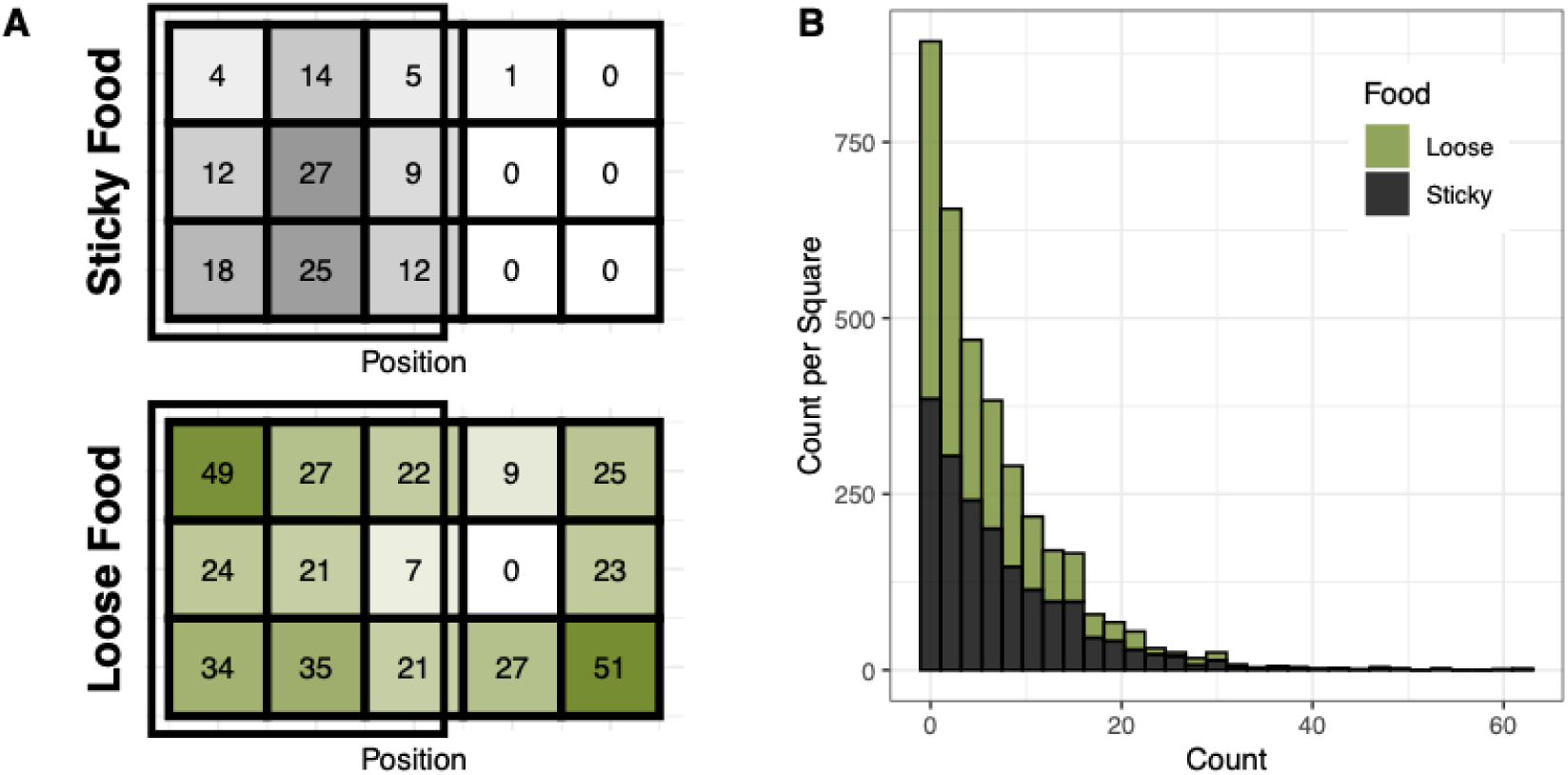
(A) Heatmaps of where individuals are found after 21 days in test microcosms of ‘sticky’ (top) and ‘loose’ (bottom) food where food containing larvae has been added to one half of the microcosm (black box) and sterile food to the other. Numbers represent the total number of individuals found in each square. (B) Histogram showing the number of grid squares (y-axis) with different counts of infected larvae (x-axis) during serial passage. Variance is higher in the ‘sticky’ food condition (MS1).

### Setting up host populations for microcosms

After thawing the pots of ‘sticky’ and ‘loose’ food, we added ∼50 similarly aged adult moths to each pot from either our Genotype 9 or Genotype 17 host maintenance populations and incubated them for 10 days. These single-genotype adults mated and laid eggs into the ‘sticky’ or ‘loose’ food. 10 days after adding adults, these pots were used to set up microcosms as in Figure 1. While pots were set up with the same number of adults, they may not have contained the same number of larvae. Observationally, population sizes may have varied from a couple dozen individuals to mid-hundreds of individuals.

Microcosms were set up in 9.25” x 12.08” x 0.9” inch plastic A4 document cases (Daiso Japan, U.S.A.) that had been marked with an even 3 x 5 grid of 3.08” x 2.42” squares. We set up 8 microcosms per replicate: 1) homogeneous Genotype 9 hosts in ‘loose’ food; 2) homogeneous Genotype 17 hosts in ‘loose’ food; 3) homogeneous Genotype 9 hosts in ‘sticky’ food; 4) homogeneous Genotype 17 hosts in ‘sticky’ food; 5) heterogeneous hosts (Genotype 9 on top / Genotype 17 on bottom) in ‘loose’ food; 6) heterogeneous hosts (Genotype 17 on top / Genotype 9 on bottom) in ‘loose’ food; 7) heterogeneous hosts (Genotype 17 on top / Genotype 9 on bottom) in ‘sticky’ food ; and 8) 7) heterogeneous hosts (Genotype 9 on top / Genotype 17 on bottom) in ‘sticky’ food (Figure 1). Each microcosm therefore had a single food type (either ‘sticky’ or ‘loose’), according to its treatment.

To arrange hosts in the microcosm, we added 2 pots of the food containing 10-day old larvae to opposite vertical ends of the grid, so that food fully filled the plastic containers. For homogeneous host treatments, we added 2 pots containing larvae of the same genotype and, for heterogeneous treatments, these 2 pots contained larvae of different genotypes. Thus, homogeneous genotype microcosms had a single genotype of host throughout the grid, while heterogeneous microcosms had two genotypes separated vertically across the grid with a contact zone at the 3^rd^ row. To balance our treatments and account for potential effects due to position within the grid, we set up 2 microcosms of each of the 2 heterogeneous treatments and alternated which genotype is on ‘top’.

These microcosms were then incubated overnight, as above, so that virus could be added to them on day 11. We repeated this process each passage to set up new microcosms so that evolving virus were added to a fresh microcosm. This allowed us to maintain 2 non-interbreeding and non-evolving host genotypes in our microcosms and preserve the spatial structuring of such genotypes across passages.

### Experimental evolution

We set up 3 replicates of each of our 8 unique microcosm types (24 serially passaged microcosms total). We initiated serial passage by pipetting 5ml of 10^-2^ stock virus solution in small droplets evenly over the top of each microcosm. Each microcosm was then incubated for 20 days and then frozen overnight to kill larvae for harvesting. Passage lengths (10 days before set-up, 1 day overnight, 20 days with virus) therefore roughly correspond to the host’s natural ∼monthly cyclical demography [40]. The average ‘time to kill’ for the ancestral virus on third instar larvae is ∼15.5 days (range 6-24), so this also roughly corresponds to a natural infection cycle (Figure S2). We stagger the replicates’ passaging schedules weekly due to the time intensive nature of passaging.

To harvest virus, we dissected each microcosm by grid square and separately collected infection killed cadavers from each square in sterile 1.5ml Eppendorf tubes (Figure 1). We added 1ml sterile, MilliQ water to each square’s tube and used a sterile pellet pestle (Fisher Scientific, U.S.A.) to manually burst larvae and release viral occlusion bodies. We dropleted 500ul of this solution from each square onto the same square of that treatment’s freshly set-up microcosm to infect the next passage and froze the rest of the solution. Therefore, we preserved the force of and spatial structure of infection within the grid across passages. We recorded the number of infected cadavers in each square at each passage to track the spatial ecology of infection in the experiment.

The microcosms newly infected with passaged virus were then placed back into incubators for 20 days to allow the virus to infect larvae for the next passage. During this time, larvae disperse from their initial position at a rate determined by their food viscosity, ingest virus from the sites they move between, and carry their infection to their eventual site of death. This means that larvae in heterogeneous sticky food microcosms are both less likely to become infected by virus from the alternate host type and are less likely to move across the contact zone to infect the alternate host type. We experimentally evolved virus for 10 passages, a number standard for microbial evolution experiments.

There were several occasions where we had to use the frozen virus solutions reserved during passaging to re-set up passages due to low numbers of infected cadavers or contamination with another pathogen. Low infected numbers were caused by low total population sizes and tended to happen in batches when a set of adults did not breed for whatever reason. There were no observable trends in when this would occur. Bt contamination was much rarer and also tended to happen in batches (potentially when a food batch had been contaminated) without observable trends. Contamination was recognizable because the most common contaminant, *Bacillus thuringiensis* (Bt), turns infected larvae black rather than white [49]. We could generally expect any co-infected larvae to display the black phenotype since Bt kills larvae more quickly (Yitbarak, unpublished data), so re-setting up passages when Bt was visually detected would clear the virus population and prevent co-infection from selecting on evolving virus. Congruously, we did not see any signs of persistent contamination within our passages nor in any of our assays.

### Assay

After passage 10, we collected virus infected cadavers from each square as above. We next pooled the virus from the 3 squares in each row of the 3x5 grid in equal proportion so that we had 5 virus samples per microcosm. Each pooled sample was therefore the virus population at a certain vertical position within the microcosm, along the gradient of heterogeneity for the heterogeneous host microcosms. We purified virus by centrifuging for 1 min at 3000 rpm to remove larger particulate matter and then 3 min at 13000 rpm to pellet virus. We ran these samples through a .65 micron filter to semi-purify our virus of larger bacterial and fungal contaminants, as in [38]. Next, we quantified the concentrations of each sample by counting occlusion bodies on a Petroff-Hauser counting chamber with 400x darkfield microscopy and diluted each to an assay dose concentration of ∼7.5x10^8 occlusion bodies per mL in 2% sucrose and .2% dye [38]. The sugar entices the larvae to ingest the virus and the dye allows the experimenter to determine which have ingested half their body lengths of solution and are considered exposed.

We assayed each pooled virus sample on both of our two host genotypes, Genotype 9 and Genotype 17, in a balanced, batched design (see Table S3) to determine the proportion of hosts each virus sample infects on each genotype and the average number of occlusion bodies each virus sample produces per infection on each genotype. With 5 virus samples for each of the 8 microcosms that have 3 replicates, this resulted in 240 virus sample x assay genotype combinations. Because only 1 genotype is assayed per day and there are strong day effects within the system, we included assay genotype and set-up date in all statistical models to account both for any general differences in resistance and assay day effects. Furthermore, conclusions are drawn from comparing the relative phenotypes of evolved virus on the different assay genotypes rather than comparing to the ancestral population that was assayed separately (Figure S3, Table S2).

To set up infectivity assays, we first moved ∼70-80 adult moths of the appropriate host genotype into new pots with 200g ‘standard’ food. Eleven days after setting up assay pots, we collected 100 third instar larvae for each assay combination in a petri dish and starved them under a damp paper towel for 2 hours. After starvation, we syringed tiny droplets of the appropriate virus-sucrose-dye solution onto the petri dish for the larvae to consume. We added 50 larvae that have orally ingested half their body lengths of virus solution to 2 25-cell compartmentalized square petri dishes (ThermoFisher Scientific, U.S.A.) with ‘standard’ food and incubated them for 20 days [38]. Assay grids were labelled with random identifiers to blind assay combinations and prevent bias. After 20 days, assay grids were frozen and destructively sampled to count the number of infected and uninfected individuals. Infected cadavers from each assay grid were then saved in a pooled sample for virus quantification.

To determine the average number of occlusion bodies produced per infected individual, we extracted virus from the pooled samples from each assay grid using a sterile pellet pestle (Fisher Scientific, U.S.A.) to manually burst larvae and release viral occlusion bodies. We then centrifuged these samples as above, but did not filter them, and counted occlusion bodies on a Petroff-Hauser counting chamber. The concentration of occlusion bodies was then divided by the number of infected cadavers that were in the pooled sample to get the average number of occlusion bodies produced per infection for each virus sample on each host genotype.

### Statistical analyses

We analyzed our data using a generalized linear mixed modelling framework in R [50]. All models were run in R version 4.2.3 (2023-03-15) -- "Shortstop Beagle" [51]. We used packages ‘glmmTMB’ and ‘lme4’ to build models, ‘DHARMa’ to check model residuals, ‘afex’ and ‘car’ to determine significant model terms, ‘emmeans’ to extract effects, ‘tidyverse’ to manipulate data, and ‘patchwork’ and ‘ggplot2’ to plot results [52–60]. We determined error structures for models by testing fitted models with ‘DHARMa’ and then adjusting to best fit residuals. On occasion, observation level random effects were used to correct overdispersion in our models [61]. Where necessary, contrasts were used with ‘emmeans’ to determine effect estimates for treatments. All model tables and annotated code are in the supplement. Models are numbered in the table (MS, M1-6) with suffixes indicating the response variable being tested (i for infectivity, p for productivity, and c for composite exploitation rate) and referenced in the manuscript as appropriate.

### Spatial structure analysis

First, we analyzed how larvae moved in our microcosm food viscosity test (Figure 2A) by asking whether the total count of individuals in a square is predicted by an interaction between the viscosity of the food and the distance (in square number) that the square is from where larvae were added (with squares where larvae were directedly added coded as 0) (MS2). The model uses a Poisson error structure with observation level random effects.

We next examined how food viscosity spatially structured infection during the evolution experiment using the data collected on the number of infected individuals in each square at each passage for each microcosm. To do this, we ran a Levene’s Test for Homogeneity of Variance on the infected count data to determine if our ‘sticky’ and ‘loose’ food treatments differ in their variance, a measure of how ‘clumpy’ infection is within a microcosm (MS1).

### Assay analysis

Infectivity and productivity assay data were analyzed in a generalized linear mixed modeling framework using the same packages and process as above [50]. We built models treating proportion infected, average number of occlusion bodies per infected individual, and the composite exploitation rate (proportion infected multiplied by the average virus count) as our response variables. We transformed average virus counts and composite exploitation rates by multiplying and rounding them to produce integer counts for Poisson and negative binomial distribution assumptions. In each model, we included random effects for batch and the nested effect of virus sample under replicate under treatment.

First, we asked whether there are significant interaction effects between the genotype that the virus is being assayed on, the host treatment that the virus is from, and the spatial structure that the virus is from by building models with a three-way interaction between these terms (M1). To more closely look for trends in our data, we asked whether virus populations significantly differ in their specialization at the whole microcosm level by building models that include an interaction effect between the assay being on the host that the population was evolved on (‘familiar’) or the foreign host and the type of food the population evolved in (M2). Virus from heterogeneous populations was coded as ‘heterogeneous’ as they do not have a familiar or foreign host at the whole microcosm level. We also included assay genotype as a fixed effect to account for any overall differences in resistance between our assay host genotypes. Model convergence issues prevented us from including the evolution genotype in all models, but, since this term had very weak effects (p>0.9) in the models that did converge, we dropped the term to improve comparability across models. The infectivity model uses a binomial error structure with observation level random effects, the productivity model uses a Poisson error structure with observation level random effects, and the exploitation rate model uses a zero-inflated negative binomial error structure with observation level random effects. To better see the effects of specialization in the homogeneous populations, we also ran these same models for the homogeneous host treatments only with the infectivity model using a binomial error structure, the productivity model using a Poisson error structure, and the exploitation rate model using a negative binomial error structure with observation level random effects (M3).

Next, we asked whether spatial structure or host heterogeneity have significant effects on viral phenotypes across genotypes (without accounting an effect of specialization). We built models for infectivity, productivity, and composite exploitation with the same random effects as above, and without fixed effects for being assayed on the familiar or foreign host. For host heterogeneity models, we included the assay genotype, food type, and whether the treatment had homogeneous or heterogeneous hosts as fixed effects, allowing for an interaction between food type and treatment heterogeneity (M4). The infectivity model uses a binomial error structure with observation level random effects and the productivity and composite exploitation models use a zero-inflated negative binomial error structure with observation level random effects. For spatial structure models, we include the food type and assay genotype as fixed effects with the same random effects as above (M5). The infectivity model uses a binomial error structure, and the productivity and composite exploitation models use a zero inflated negative binomial structure with observation level random effects.

Finally, we asked whether virus phenotypes are spatially structured within the heterogeneous host treatments (M6). First, the local proportion of host genotypes during evolution was determined for each virus population by its distance from the host contact zone so that the edge of the microcosm where host genotype 17 was added was coded as 1, the next row of the grid was coded .75, the middle contact zone was coded as 0.5, the following row was coded as .25, and the edge of the microcosm where host genotype 9 was added was coded as 0. Thus, the inverse represents the local proportion of host genotype 9. We built models for infectivity, productivity, and exploitation rate that included the local proportion of host genotype, the host genotype of the assay, the food type, an interaction effects between these three terms, and the order of genotypes in the microcosm as fixed effects. We included the same random effects as in the previous models for batch and nestedness and also for the specific position. The infectivity model uses a binomial error structure, the productivity model uses a Poisson error structure with observation level random effects, and the exploitation rate model uses a zero-inflated negative binomial error structure.

## RESULTS

### Food viscosity spatially structures infection

In our experiments testing how food viscosity affected larval movement, we see that ‘sticky’ food greatly decreases larval dispersal (Figure 1A, Table S1). In test microcosms where larvae are only added to one side, the count of individuals found in a grid square after 21 days i significantly affected by the interaction between the distance that the square is from where larvae were added and the food viscosity (MS2 ANOVA p < 0.001), with larvae in sticky food dispersing less (MS2 GLMM effect estimate = -3.8, p < 0.001). Interestingly, neither the effect of distance (p = 0.96) nor food viscosity (p = 0.07) significantly affects the count in a grid square alone (MS2 ANOVA).

Across the evolution experiment, microcosms had an average of 199.7 infected cadavers (range 23-626). Treatments with different food viscosities differ in the variance of number of infected cadavers per square (p < 0.001, MS1), with ‘sticky’ food having more clustered infected individuals, indicating more local transmission (Figure 2B).

### The impact of specialization on virus phenotypes depends on spatial structure

Without allowing for interaction effects, neither food viscosity (M2c ANOVA p = 0.8, M2i ANOVA p = 0.7, M2p ANOVA p = 0.8) nor being assayed on the familiar or foreign host (M2c ANOVA p = 0.2, M2i ANOVA p = 0.059, M2p ANOVA p = 0.7) significantly affect evolved viru phenotypes in homogeneous microcosms (Figure S5). This is in contrast to previous results from the system that manipulate spatial structure, but only assay on familiar, genetically diverse hosts [30] and that assay on both familiar and foreign hosts, but do not manipulate spatial structure [38].

However, trade-offs associated with virus host range and virus life history strategy may interact to change evolutionarily optimal strategies for a population. When we build models that allow for interactions between spatial structure and host specialization, we see that the effect of being assayed on the familiar line depends on spatial structure (M3c ANOVA p = 0.047, GLMM effect estimate Foreign x Sticky = 0.10 p = 0.047) (Figure 3, S4).

**Figure 3:**
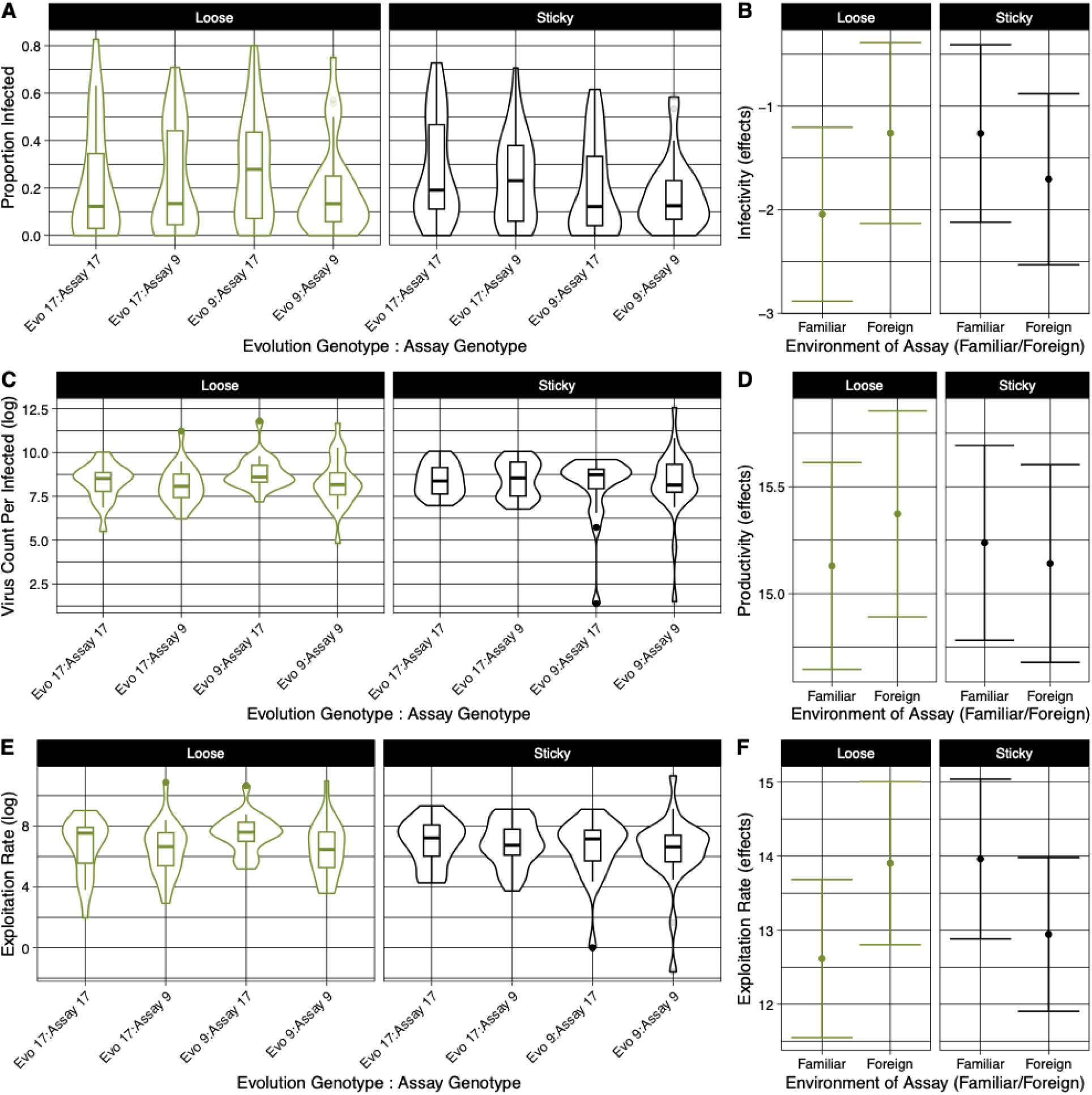
Interactions between specialization and the spatial structuring of infection in homogeneous microcosms. Panels A, C, E plot raw data and panels B, D, F show model effect estimates drawn from M3 GLMMs. Y-axes represent: (A) proportion infected, (B) effect estimates for the M3i infectivity model, (C) average number of occlusion bodies produced per infectious cadaver on a log scale, (D) effect estimates for the M3p viral productivity model, (E) composite exploitation rate (proportion infected * average number of occlusion bodies) on a log scale, and (F) effect estimates for the M3c exploitation rate model. A, C, E X-axes represent the combinations of whic host genotype the virus population evolved on and which they were assayed on. B, D, F X-axes represent whether the virus was being assayed on the host genotype that it evolved on (familiar) or on the foreign host genotype. Panels show whether the virus came from a treatment with ‘loose’ food or ‘sticky’ food.

Virus populations are significantly less exploitative of their familiar host in loose food (M3c contrasts loose effect estimate = -1.29 p = 0.03) and non-significantly more exploitative of their familiar host in sticky food (M3c contrasts sticky effect estimate = 1.02 p = 0.09) (Figure 3F). This is largely because of differences in viral productivity (M3p contrasts loose effect estimate = -0.78 p = 0.03 ; sticky effect estimate = 0.442 p = 0.2) (Figure 3D), rather than infectivity (M3i contrasts loose effect estimate = -0.24 p = 0.37 ; sticky effect estimate = 0.09 p = 0.7) (Figure 3B).

This indicates that the direction of the effect of viral specialization on host exploitation rate varies with population spatial structure.

### Spatially structured host genetic diversity leads to local specialization

On average, virus populations evolved in heterogeneous host populations do not significantly differ in infectivity (M4i ANOVA, p = 0.8) or composite exploitation (M4c ANOVA, p=0.12) from those evolved in homogeneous host populations and are slightly more productive (M4p GLMM effect estimate = 0.16 p = 0.04).

However, heterogeneous host populations have gradients of host mixing within them as the different positions are different distances from the other host genotype. Additionally, we expect spatial structure to impede host movement and thus further alter the degree of host mixing for different positions. Therefore, we explore whether there is an effect of host specialization within the heterogeneous host microcosms and whether this effect is altered by the spatial structuring of the population.

We find that, for composite exploitation rate, there is a significant interaction between the local proportion of a host genotype during evolution (i.e. the position’s distance from the contact zone), the assay host genotype, and the type of food evolved in (M6c ANOVA p = 0.029, GLMM estimate = 0.56 p = 0.029, Figure 4). This is due to differences in virus productivity (M6p ANOVA p = 0.018), but not infectivity (M6i ANOVA p = 0.60).

**Figure 4:**
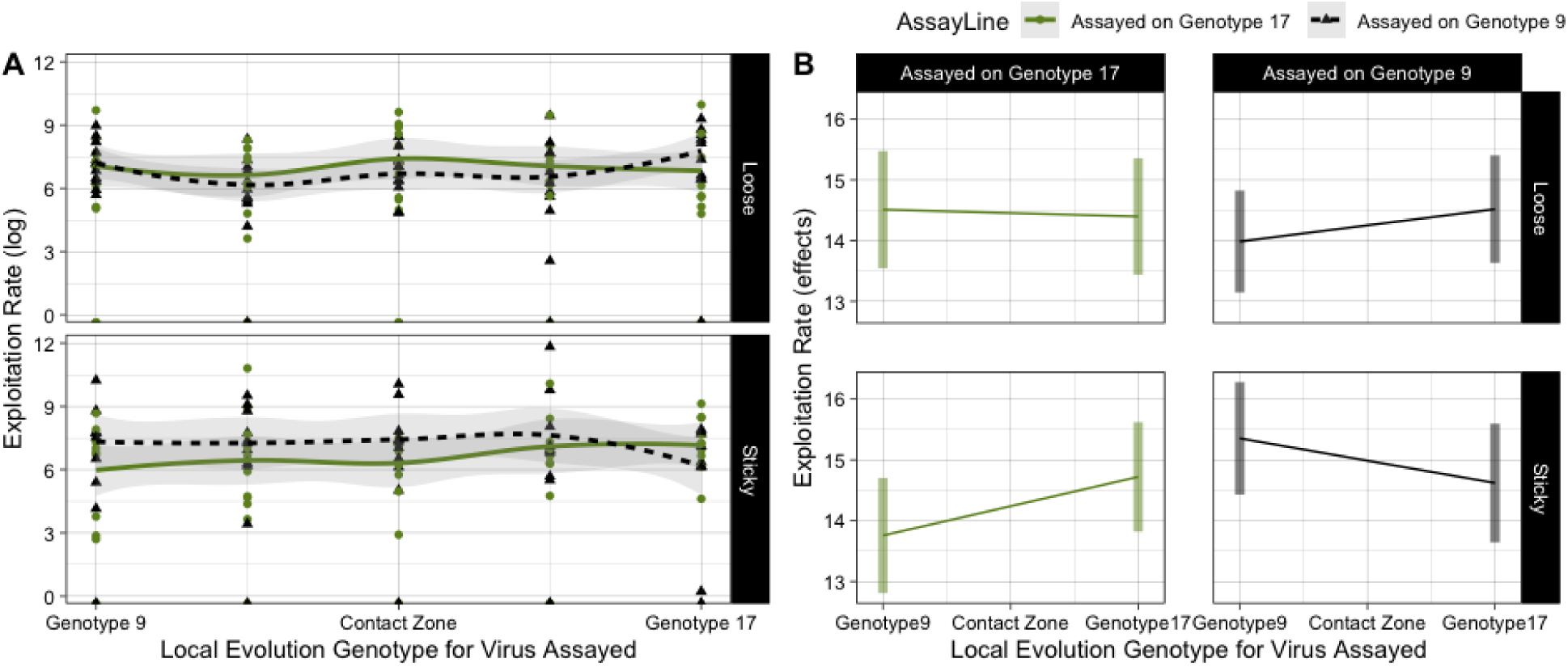
Spatially structured host heterogeneity results in locally specialized virus. Virus sampled from different spatial positions within each microcosm (along x-axis with tick labels denoting the local evolution host genotype, i.e. Figure 1 vertical axes) were assayed on the two host genotypes. (A) Raw data for the composite exploitation rate (proportion infected * average number of occlusion bodies, log-scale, y-axis) on the two host genotypes (green, circle, solid line is exploitation rate on genotype 17; black, triangle, dashed is on genotype 9) for virus from ‘sticky’ and ‘loose’ food microcosms (panels) for virus sampled from different spatial positions within the microcosm (x-axis) with therefore different evolutionary histories of mixing between the 2 host genotypes (B) Model effect estimates drawn from M6c GLMM for the interacting effects of food type and assay genotype on exploitation rate (y-axis) for virus sampled from different spatial positions within the microcosm with different degrees of mixing between the 2 host genotypes (x-axis). Panels are separated vertically by food type and horizontally by which host genotype the virus was assayed on.

The significant interaction between local proportion of a host genotype during evolution, assay host genotype, and food type is driven by populations from sticky food where virus from different positions within the microcosm have higher exploitation rates on their locally familiar host populations (M6c pairwise contrasts effect estimate = 1.7 p = 0.033). In contrast, viru phenotypes within loose food microcosms do not significantly vary, though, like in homogeneous microcosms, they have slightly lower exploitation rates on the familiar host (M6c pairwise contrasts effect estimate = -0.65 p = 0.37). This indicates that spatially structured host heterogeneity results in the evolution of more diverse, locally specialized virus.

## DISCUSSION

We experimentally evolve baculovirus in microcosms that vary in spatial structure and host genetic diversity to explore how these two selection pressures interact to shape host range evolution. Our experimental evolution approach manipulates the viscosity of the food medium that the moth larvae live within and adds either homogeneous (single inbred line) or heterogeneous (two inbred line) hosts. We then assay virus population from different positions within the microcosm on both the familiar and foreign host. The key result of our experiment is that spatially structured heterogeneous host populations support localized virus specialization and, therefore, more diverse viral strategies within the population. Virus sampled from different positions within heterogeneous microcosms showed higher exploitation rates on the locally dominant host genotype in spatially structured environments but not in well-mixed ones (Figure 4). This finding supports theory suggesting that more contact between host genotypes selects for generalism, while lower contact rates can allow for the persistence of local specialists [20,22]. This implies that the benefits of host genetic diversity in selecting for ‘mediocre generalists’ and preventing the ‘monoculture effect’ [2,4,19] may only be realized when host heterogeneity is well-mixed.

A second important result is that we also find that the direction of the effect of specialization on host exploitation rate is reversed depending on spatial structure. Specifically, in both homogeneous and heterogeneous treatments, virus populations had higher exploitation rates on their familiar host in spatially structured microcosms, and lower exploitation rates on their familiar host in well-mixed microcosms (Figure 3). This suggests that spatial structure may be imposing additional selection pressures that are altering the evolutionarily optimal strategy on the familiar host type. However, it can be argued that the direction of the specialization results is counterintuitive. Standard expectations—based on earlier theory and experiments—would predict that well-mixed microcosms should select for increased exploitation of the familiar host, while spatially structured environments might select for more prudent exploitation [27,28,30]. We saw the opposite.

There are several potential explanations for this unexpected result. The first is that some theory suggests that intermediate levels of spatial structure can actually select for higher parasite exploitation rates [28,33]—our sticky food treatment could fall within this range. However, this would not explain why viruses from well-mixed treatments evolved lower exploitation rates on their familiar host. These lower exploitation rates could, however, result from an unmeasured phenotype, such as time to kill, being selected upon and trading off with our measured metrics (infectivity and productivity). In other baculoviruses, shorter time to kill correlates with lower virus productivity [62], so selection for rapid infection cycles could explain the lower exploitation rates we observe in loose food conditions. In addition, another phenotype that we did not measure but that could trade-off with viral infectivity is environmental persistence [63]. Finally, it could also be possible that the loose food environment more closely resembled ancestral conditions, resulting in weak selection on exploitation rate, while virus in sticky food experienced stronger selection to increase exploitation on the familiar host. Meanwhile, exploitation of the foreign host may have drifted in opposite directions independently.

Our results have implications for broader ecological and evolutionary theory on how specialization supports diversity, as they challenge the general assumption of rare host advantage. In our experiment, pathogens sometimes exhibited (presumably maladaptive) high virulence on foreign hosts, suggesting that rare host genotypes may not always have a fitness advantage. This would complicate many models of host–parasite coevolution—including the Red Queen Hypothesis and Janzen-Connell effects [64–66]—that depend on negative frequency-dependent selection arising from rare host advantage. We find that, under certain spatial and population structures, selection for lower virulence on the familiar host type can make rare hosts more vulnerable to infection. While similar patterns of high foreign host virulence have been noted in contexts such as species invasions (i.e., "invading with weapons") and zoonoses [67,68], those cases are typically attributed to evolved resistance in the familiar host leading to coevolutionary arms races that result in higher foreign host virulence. Our results suggest that such patterns can occur even in the absence of host coevolution. If such reversed specialization emerges regularly under certain ecological conditions, it may violate the assumptions necessary for rare genotype advantage to maintain diversity through negative frequency-dependent selection.

In conclusion, our experiment shows that spatial structure and host genetic diversity interact to alter pathogen host range evolution and how pathogens specialize. We demonstrate that spatially structured host heterogeneity can support local specialization, and that the direction of specialization’s effect on exploitation rate—whether higher or lower on the familiar host— depends on the population’s spatial structure. These findings highlight the importance of multidimensional trade-offs in pathogen evolution, particularly interactions between host range and life-history traits, and suggest that such interactions can produce unexpected and context-dependent outcomes. Finally, this work has important implications for controlling infectious diseases in heterogeneous populations, as spatial structure may limit the effectiveness of host diversity in constraining pathogen adaptation.

