## Supplementary material for "Spatially structured host genetic diversity leads to the evolution of local specialization": Figure S , Table S

**Figure S1:** Box plots illustrating life‐history traits (growth rate, development time, pupal mass) of Inbred Line 9 & 17 for different glycerol levels: 35mL Gly (Food 1), 48.75mL Gly (Food 2), 62.5mL Gly (Food 3), 76.25mL Gly (Food 4), 90mL Gly (Food 5). All food was made with Earth’s Best breakfast cereal (100g), brewer’s yeast (20g), sorbic acid (0.2g), and methyl paraben (0.2g). Line and Food type do not significantly affect key life history metrics.

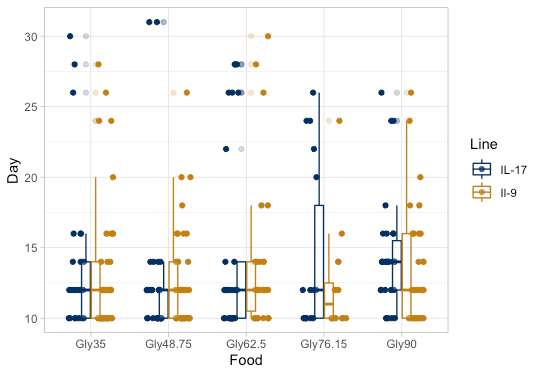

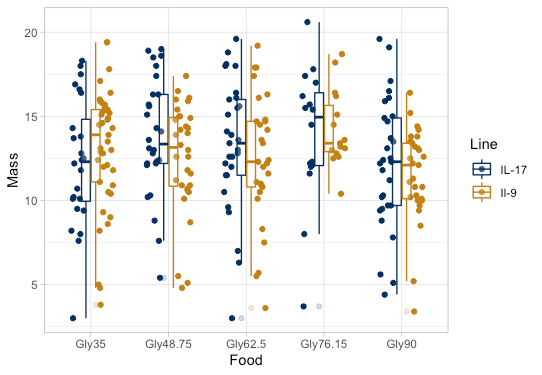

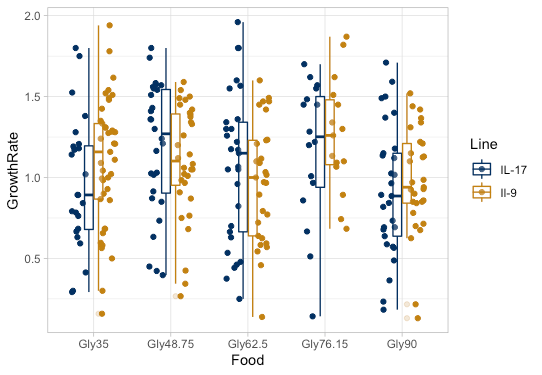

Growth Rate (mass /days)

Mass at pupation

Days until pupation

**Table S1:** Summary statistics (average, standard deviation, standard error) of IL-9 larvae movement data in spatial structure assay. Spatial assays are set up with 18cm PVC tubes with a diameter of 4cm. Both ends of the tube are double layered with mesh cloth and rubber bands to ensure that food and larvae don’t escape. Food is lightly packed into the tube by knocking it against a surface and third instar larvae are placed into one end of the tube clearly marked as the starting point. After incubating for 10-14 days, the pipes are frozen to ensure that larvae no longer move after the allotted amount of time. Frozen food is then pushed out using a flat surface to not disturb the larvae location. Starting from the larvae starting point, the food is sifted through 0.5cm at a time until the larvae is uncovered, and distance traveled by the larvae is recorded.

|  | **Loose Food** | **Sticky Food** |
| --- | --- | --- |
| **Sample Size** | 9 | 10 |
| **Average (cm)** | 2.82 | 1.77 |
| **Standard Deviation (cm)** | 3.47 | 2.00 |
| **Standard Error (cm)** | 1.16 | 0.56 |

**Figure S2:** Time to kill for the ancestral virus when infecting 3^rd^ instar larvae. Ancestral virus was assayed on 3^rd^ instar larvae from inbred line 10 and plates were checked daily post infection. Time of death (when the larvae turned opaque white and stopped responding to stimulus) was recorded. Average time to kill was 15.5 days and ranged between 6-24.

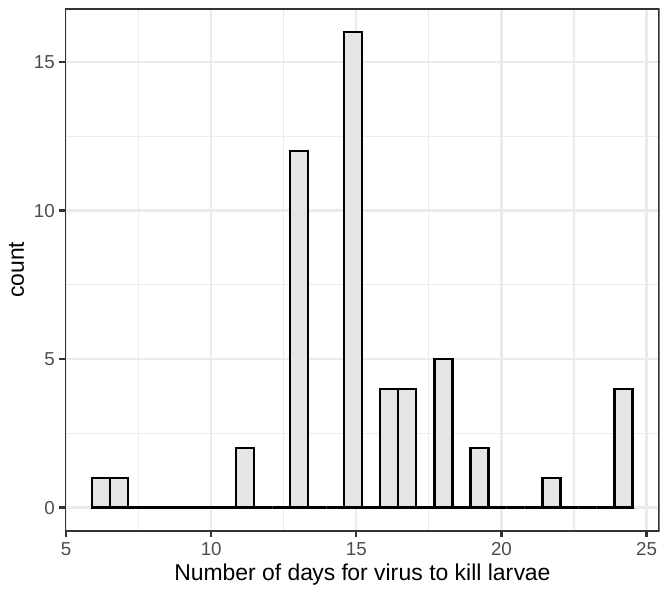

**Figure S3:** Raw data divided by treatment (x-axis) for: (y-axes) (A) proportion infected, (B) average virus count per infected cadaver (log scale), and (C) composite exploitation rate (log scale. Plots are faceted by the host genotype that the assay was conducted on. Horizontal red dashed lines mark ‘ancestral’ phenotypes calculated from Visher et al. (2022) by averaging passage 1 virus lines’ phenotypes on each assay line. P1 is chosen (rather than P0 (shown in Table S2)) because differences in virus purification methods between stock, ancestral virus and evolution experiment virus (used in Visher et al. (2022) and these experiments) led to differences in per particle infectivity. Averaged P1 metrics should approximate starting phenotypes because little evolution had occurred by P1 in Visher et al. (2022) and we average the metrics of lines whose single passage was on the assay line and lines that were on alternate host lines. Note, however, that assays on genotype 17 and on genotype 9 were conducted on different, single days and so likely have strong batch effects between them.

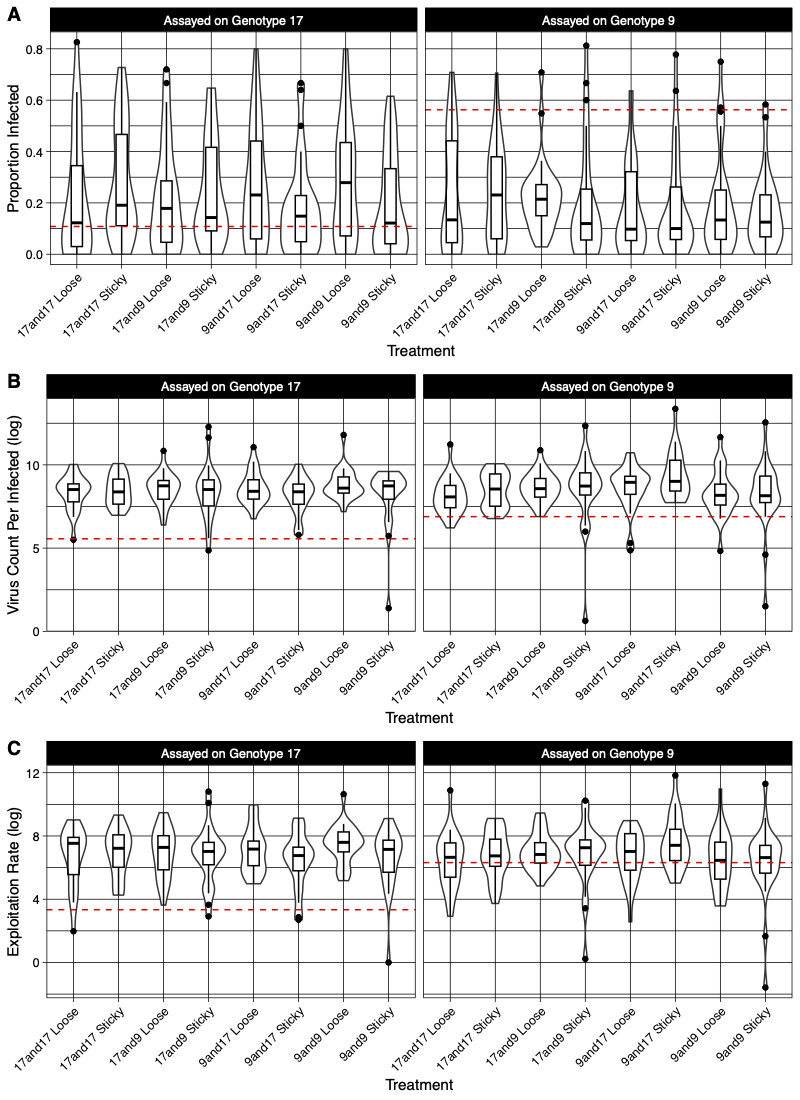

**Figure S4:** Model effect estimates drawn from GLMMS for the interaction between assay genotype : evolution genotype(s) : food type (x-axis) at the whole microcosm level for: (y-axes) (A) proportion infected, (B) average virus counts per infected cadaver (log scale), and (C) composite exploitation rate (log scale. Plots are faceted by the food type.

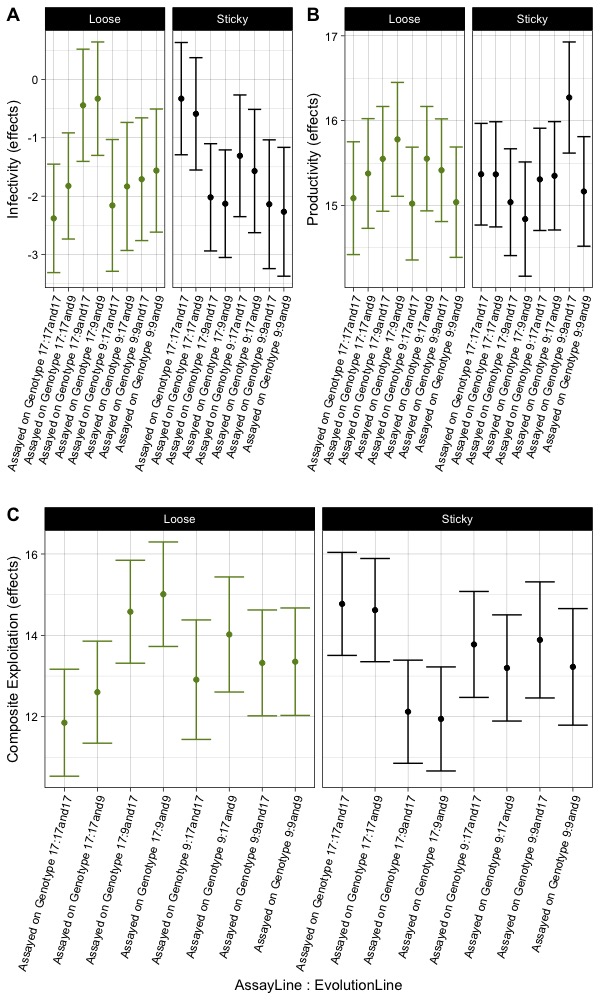

**Table S2:** Virus infectivity of ancestral virus among host genotypes (i.e. general host resistance or starting virus phenotypes). Note that these assays were conducted at different concentrations as experimental assays and differences between virus purification methods between these, and experimental virus populations, led to different per particle infectivity rates.

|  | Line 17 | Line 9 |
| --- | --- | --- |
| Proportion infected at 5.6 x 10^8 OB/mL | 7/11 | 7/9 |
| Percent | 63.6% | 77.8% |
| Proportion infected at 2.8 x10^8 OB/mL | 6/15 | 10/20 |
| Percent | 40% | 50% |

The passage dose was ~7.5x10^8 occlusion bodies per mL.

**Figure S5:** Violin plots showing the overall impact of spatial structure (x-axis) on (A) proportion infected, (B) average virus count per infected cadaver (log scale), (C) composite exploitation rate (log scale).

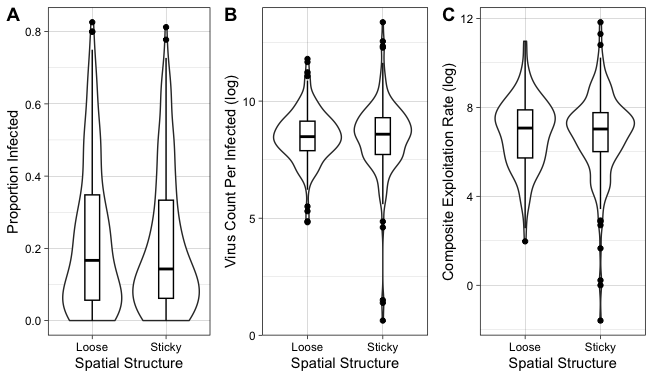

**Table S3:** Batching design for assays. One host is assayed on each batch and batches are balanced for food viscosity, lines, and heterogeneity.

| **Batch** | **Assay Genotype** | **Replicate** | **Lines** | **Space** | **Virus from Positions** |
| --- | --- | --- | --- | --- | --- |
| 1 | 17 | 1 | 17and17 | Loose | top, top middle, middle, middle bottom, bottom |
| 1 | 17 | 1 | 9and9 | Sticky | top, top middle, middle, middle bottom, bottom |
| 1 | 17 | 1 | 17and9 | Loose | top, top middle, middle, middle bottom, bottom |
| 1 | 17 | 1 | 9and17 | Sticky | top, top middle, middle, middle bottom, bottom |
| 2 | 17 | 1 | 17and17 | Sticky | top, top middle, middle, middle bottom, bottom |
| 2 | 17 | 1 | 9and9 | Loose | top, top middle, middle, middle bottom, bottom |
| 2 | 17 | 1 | 17and9 | Sticky | top, top middle, middle, middle bottom, bottom |
| 2 | 17 | 1 | 9and17 | Loose | top, top middle, middle, middle bottom, bottom |
| 3 | 9 | 1 | 17and17 | Loose | top, top middle, middle, middle bottom, bottom |
| 3 | 9 | 1 | 9and9 | Sticky | top, top middle, middle, middle bottom, bottom |
| 3 | 9 | 1 | 17and9 | Loose | top, top middle, middle, middle bottom, bottom |
| 3 | 9 | 1 | 9and17 | Sticky | top, top middle, middle, middle bottom, bottom |
| 4 | 9 | 2 | 17and17 | Sticky | top, top middle, middle, middle bottom, bottom |
| 4 | 9 | 2 | 9and9 | Loose | top, top middle, middle, middle bottom, bottom |
| 4 | 9 | 2 | 17and9 | Sticky | top, top middle, middle, middle bottom, bottom |
| 4 | 9 | 2 | 9and17 | Loose | top, top middle, middle, middle bottom, bottom |
| 5 | 17 | 2 | 17and17 | Loose | top, top middle, middle, middle bottom, bottom |
| 5 | 17 | 2 | 9and9 | Sticky | top, top middle, middle, middle bottom, bottom |
| 5 | 17 | 2 | 17and9 | Loose | top, top middle, middle, middle bottom, bottom |
| 5 | 17 | 2 | 9and17 | Sticky | top, top middle, middle, middle bottom, bottom |
| 6 | 17 | 2 | 17and17 | Sticky | top, top middle, middle, middle bottom, bottom |
| 6 | 17 | 2 | 9and9 | Loose | top, top middle, middle, middle bottom, bottom |
| 6 | 17 | 2 | 17and9 | Sticky | top, top middle, middle, middle bottom, bottom |
| 6 | 17 | 2 | 9and17 | Loose | top, top middle, middle, middle bottom, bottom |
| 7 | 9 | 2 | 17and17 | Loose | top, top middle, middle, middle bottom, bottom |
| 7 | 9 | 2 | 9and9 | Sticky | top, top middle, middle, middle bottom, bottom |
| 7 | 9 | 2 | 17and9 | Loose | top, top middle, middle, middle bottom, bottom |
| 7 | 9 | 2 | 9and17 | Sticky | top, top middle, middle, middle bottom, bottom |
| 8 | 9 | 2 | 17and17 | Sticky | top, top middle, middle, middle bottom, bottom |
| 8 | 9 | 2 | 9and9 | Loose | top, top middle, middle, middle bottom, bottom |
| 8 | 9 | 2 | 17and9 | Sticky | top, top middle, middle, middle bottom, bottom |
| 8 | 9 | 2 | 9and17 | Loose | top, top middle, middle, middle bottom, bottom |
| 9 | 17 | 3 | 17and17 | Loose | top, top middle, middle, middle bottom, bottom |
| 9 | 17 | 3 | 9and9 | Sticky | top, top middle, middle, middle bottom, bottom |
| 9 | 17 | 3 | 17and9 | Loose | top, top middle, middle, middle bottom, bottom |
| 9 | 17 | 3 | 9and17 | Sticky | top, top middle, middle, middle bottom, bottom |
| 10 | 17 | 3 | 17and17 | Sticky | top, top middle, middle, middle bottom, bottom |
| 10 | 17 | 3 | 9and9 | Loose | top, top middle, middle, middle bottom, bottom |
| 10 | 17 | 3 | 17and9 | Sticky | top, top middle, middle, middle bottom, bottom |
| 10 | 17 | 3 | 9and17 | Loose | top, top middle, middle, middle bottom, bottom |
| 11 | 9 | 3 | 17and17 | Loose | top, top middle, middle, middle bottom, bottom |
| 11 | 9 | 3 | 9and9 | Sticky | top, top middle, middle, middle bottom, bottom |
| 11 | 9 | 3 | 17and9 | Loose | top, top middle, middle, middle bottom, bottom |
| 11 | 9 | 3 | 9and17 | Sticky | top, top middle, middle, middle bottom, bottom |
| 12 | 9 | 3 | 17and17 | Sticky | top, top middle, middle, middle bottom, bottom |
| 12 | 9 | 3 | 9and9 | Loose | top, top middle, middle, middle bottom, bottom |
| 12 | 9 | 3 | 17and9 | Sticky | top, top middle, middle, middle bottom, bottom |
| 12 | 9 | 3 | 9and17 | Loose | top, top middle, middle, middle bottom, bottom |
