## Supplementary material for "Spatially structured host genetic diversity leads to the evolution of local specialization": M

**Model S1: Which food type spatially structured infection more during serial passage?**

leveneTest(Count~Food, long2)

*data = infection counts during serial passage experiment*

**Levene's Test for Homogeneity of Variance (center = median)**

Df F value	Pr(>F)

group	1 32.838 1.084e-08 ***

3581

---

Signif. codes: 0 ‘***’ 0.001 ‘**’ 0.01 ‘*’ 0.05 ‘.’ 0.1 ‘ ’ 1

| **Model S2: Which food type spatially structured infection more during microcosm dispersal food test?** | | |
| --- | --- | --- |
| Total~Distance*Food+(1\|ObsID)  *family =poisson*  *package = lme4*  *data = larval counts viscosity test adding larvae to only one half of microcosm (Figure 2C)* | | |
| **ANOVA** | **General Linear Mixed Model** | **GLMM Estimate Forest Plot** |
| 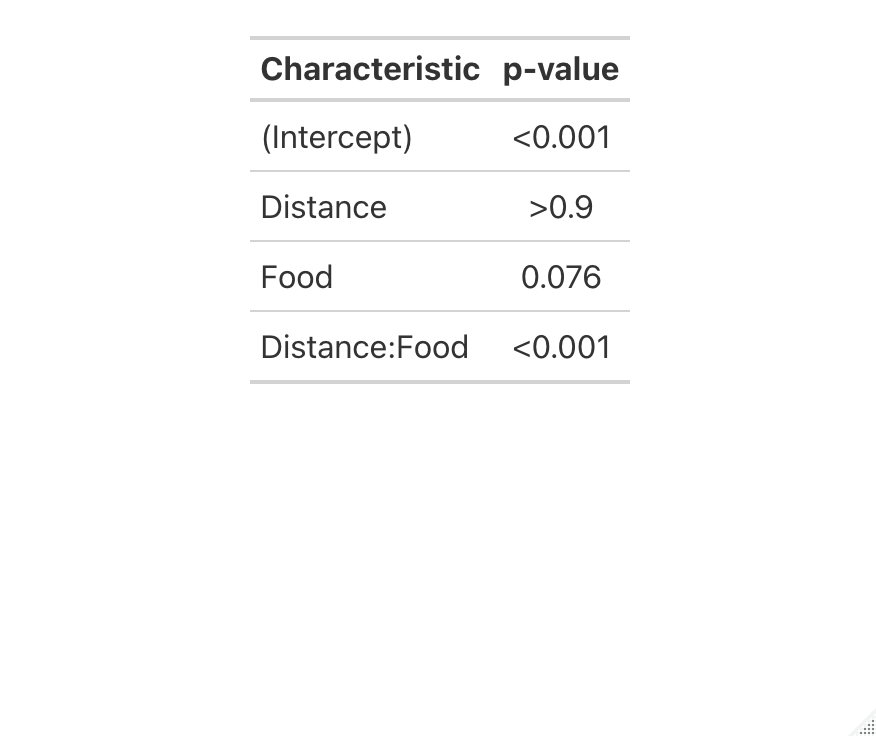 | 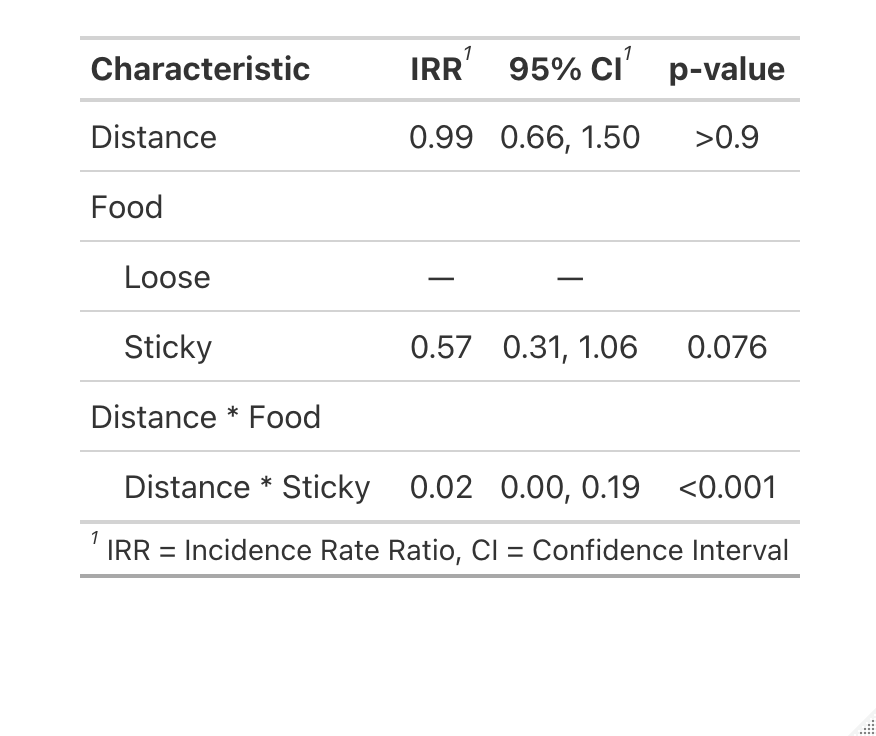 | 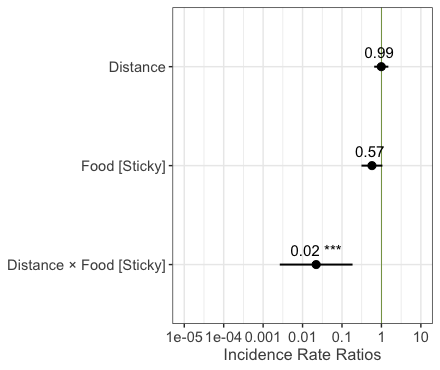 |

| **Model 1: Did assay line, evolution line(s), and/or food viscosity interact to affect virus evolution?** | | |
| --- | --- | --- |
| *Data = End of Evolution Assays* | | |
| **Composite Exploitation Rate (c)** | **Proportion Infected (i)** | **Per Infection Productivity (p)** |
| FitnessTrans ~ AssayLine * Lines * Food + (1\|Treatment/Replicate/VirusLine) + (1\|Set.Up) + (1\|ObsID)  *family = nbinom1*  *package = glmmTMB* | cbind(Infected, Uninfected) ~ AssayLine * Lines * Food + (1\|Treatment/Replicate/VirusLine) + (1\|Set.Up) + (1\|ObsID)  *family = binomial*  *package = glmmTMB* | CountPerTrans ~ AssayLine * Lines * Food + (1\|Treatment/Replicate/VirusLine) + (1\|Set.Up) + (1\|ObsID)  *family = Poisson*  *package = glmer* |
| **ANOVA**  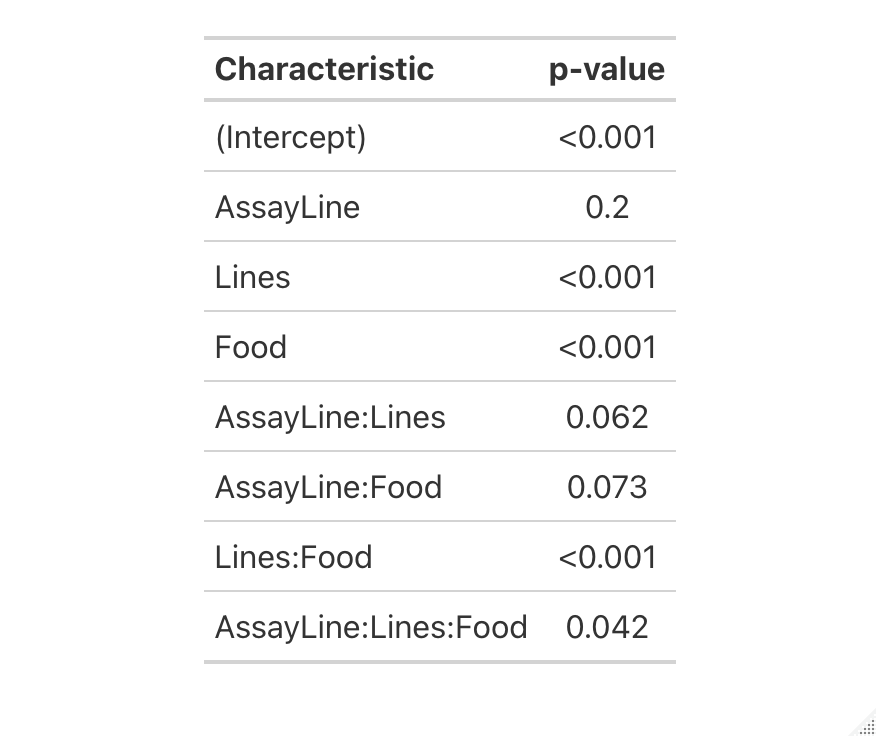 | **ANOVA**  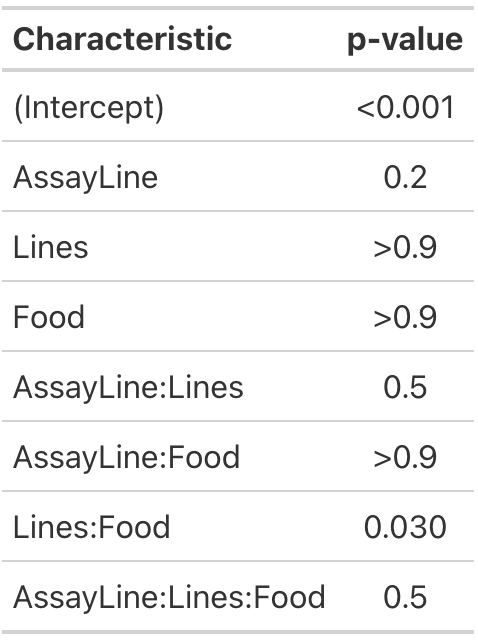 | **ANOVA**  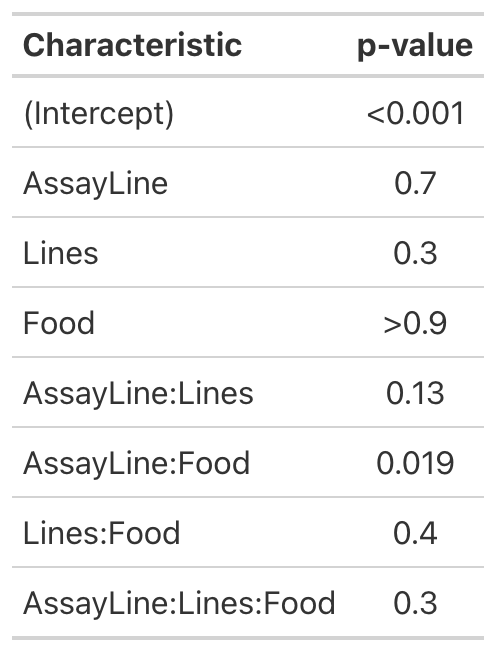 |
| **GLMM Summary**  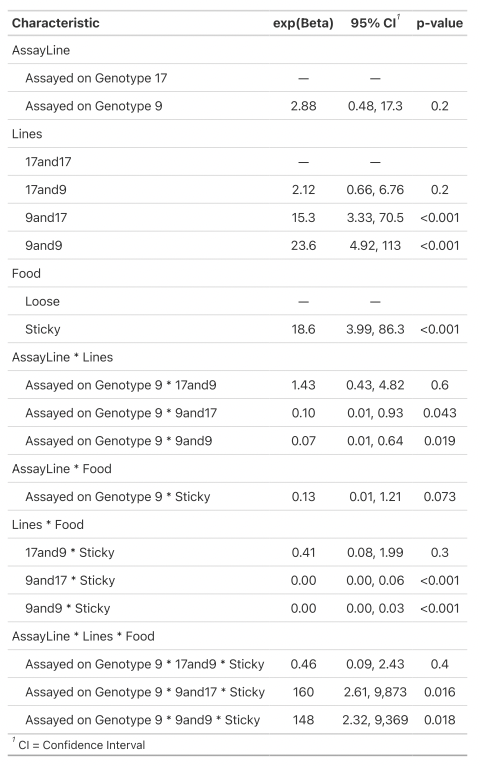 | **GLMM Summary**  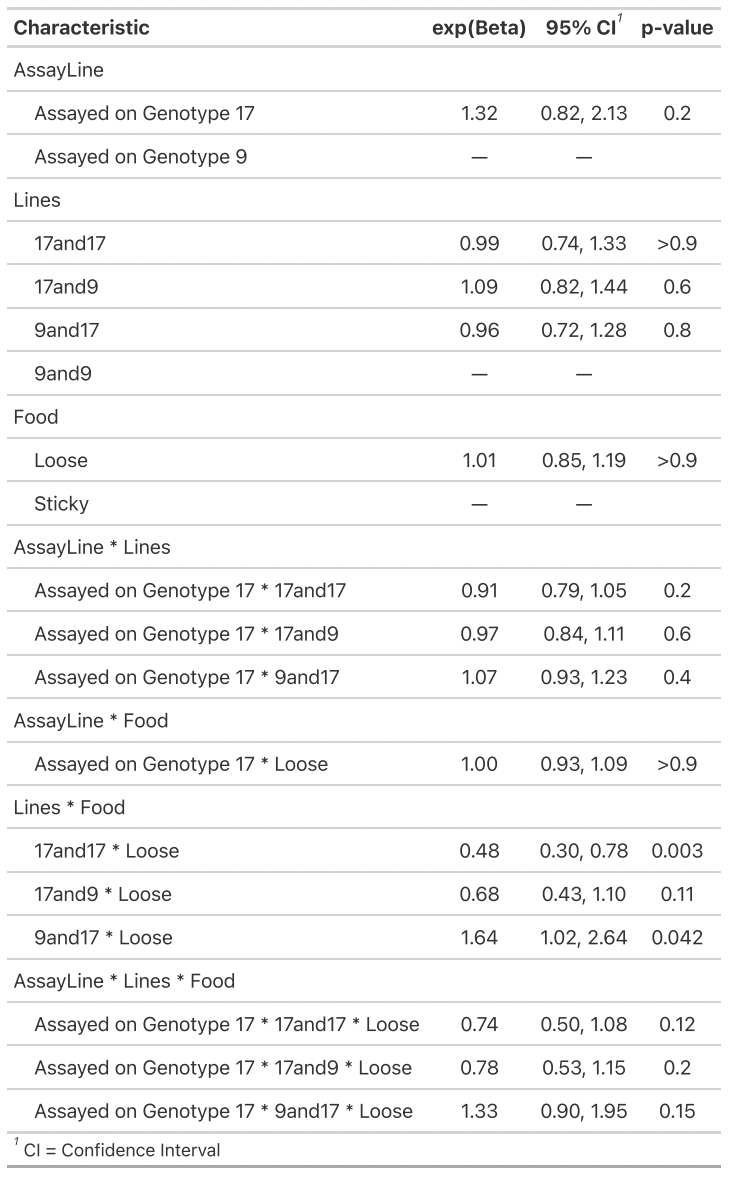 | **GLMM Summary**  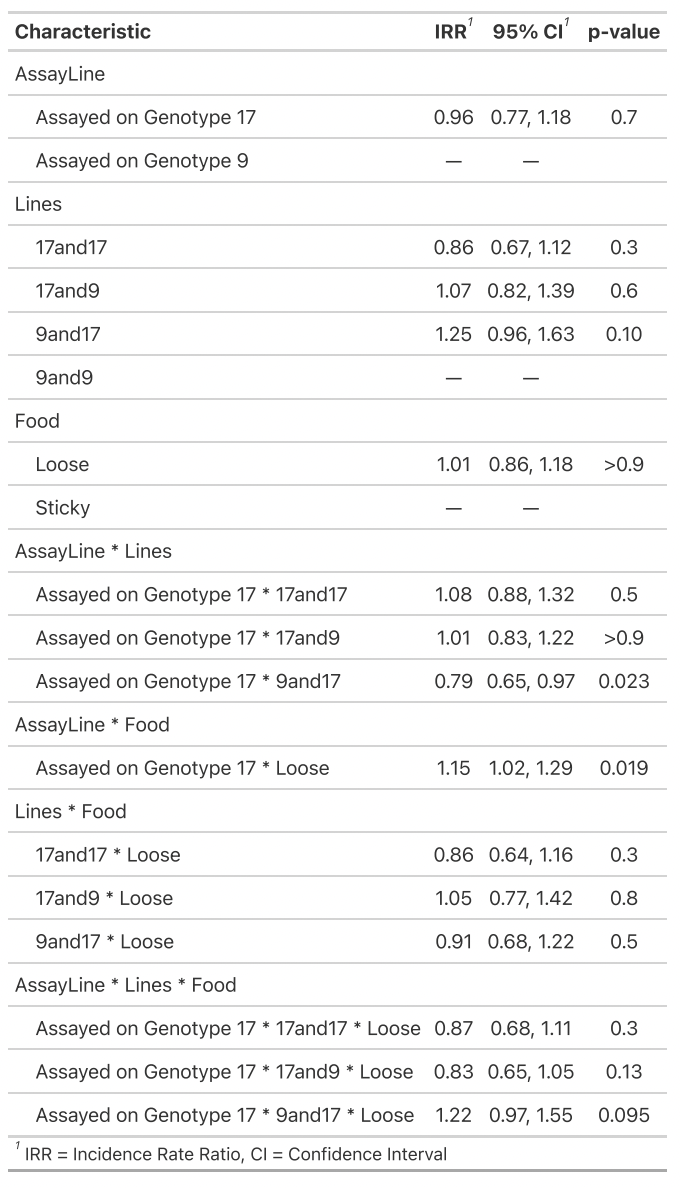 |

| **Model 2: Do either food viscosity or familiar/foreign host affect virus phenotypes w/o interactions?** | | |
| --- | --- | --- |
| *Data = End of evolution assays for homogeneous microcosms only* | | |
| **Composite Exploitation Rate (c)** | **Proportion Infected (i)** | **Per Infection Productivity (p)** |
| FitnessTrans ~ Self + AssayLine + Food + (1\|Treatment/Replicate/VirusLine) + (1\|Set.Up), zi=~AssayLine  *family = nbinom2*  *package = glmmTMB* | cbind(Infected, Uninfected) ~ Self + AssayLine + Food + (1\|Treatment/Replicate/VirusLine) + (1\|Set.Up)  *family = binomial*  *package = glmer* | CountPerTrans ~ Self + AssayLine + Food + (1\|Treatment/Replicate/VirusLine) + (1\|Set.Up) + (1\|ObsID)  *family = Poisson*  *package = glmer* |
| **ANOVA**  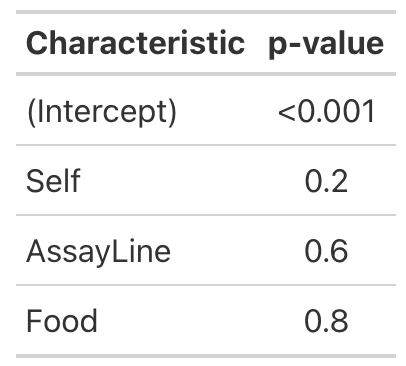 | **ANOVA**  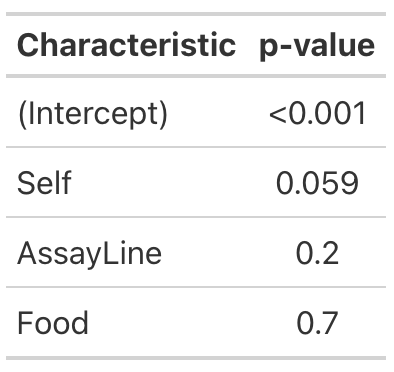 | **ANOVA**  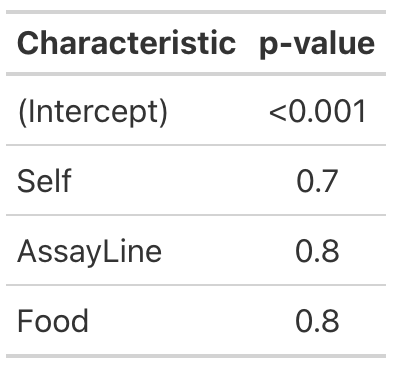 |
| **GLMM Summary**  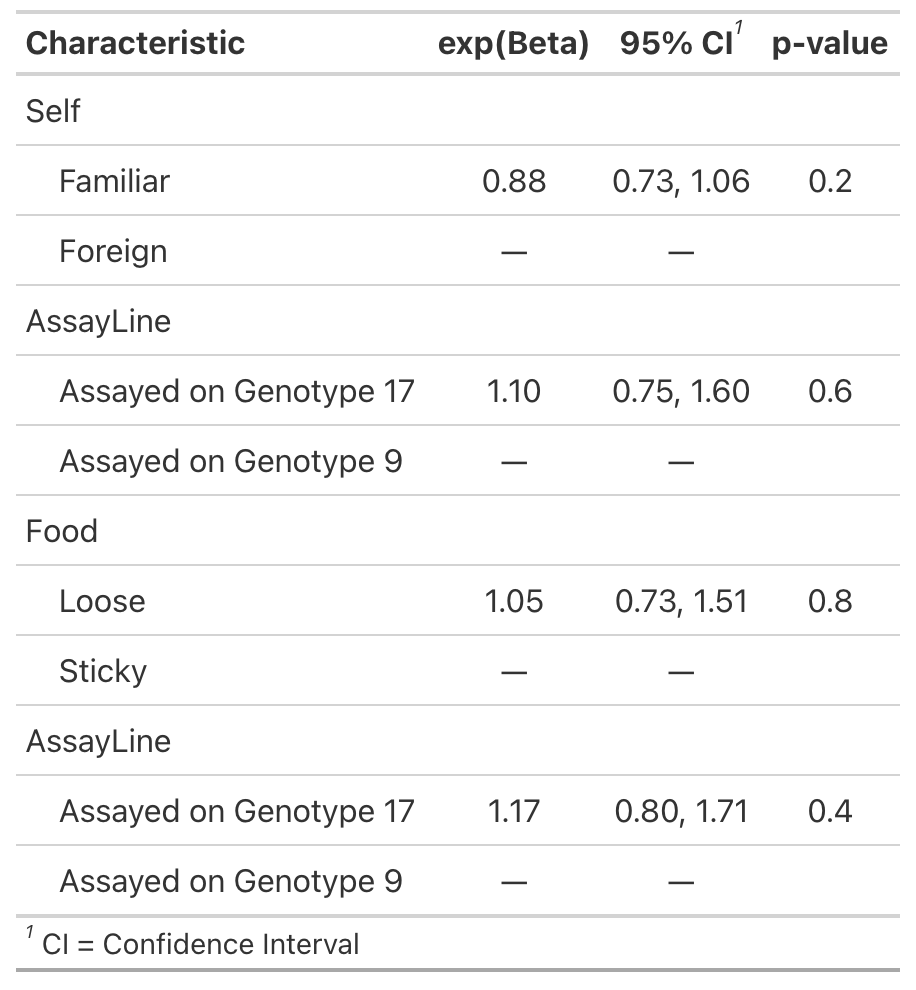  Zero Inflation | **GLMM Summary**  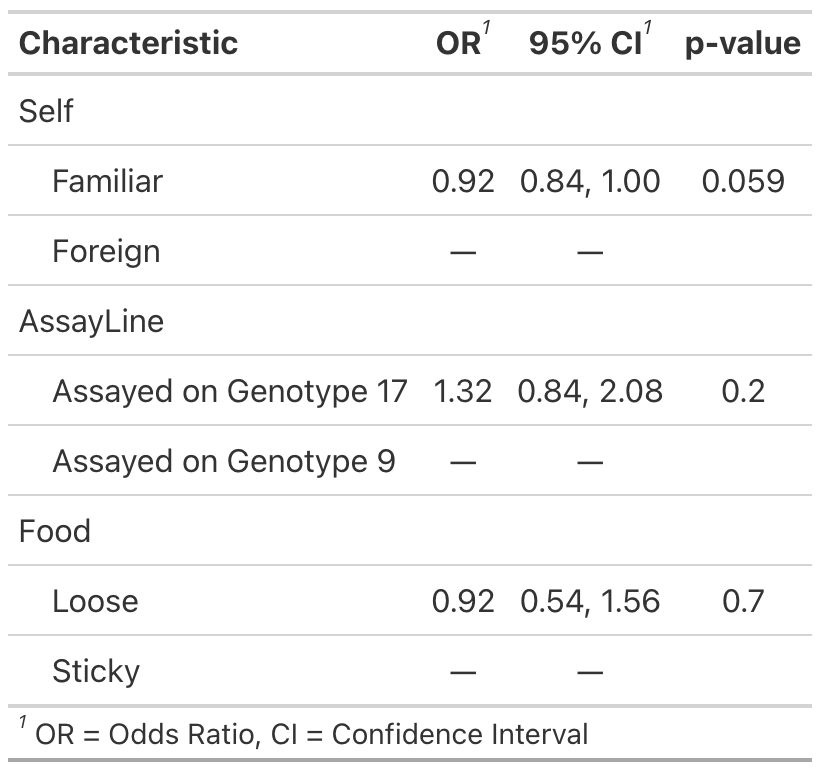 | **GLMM Summary**  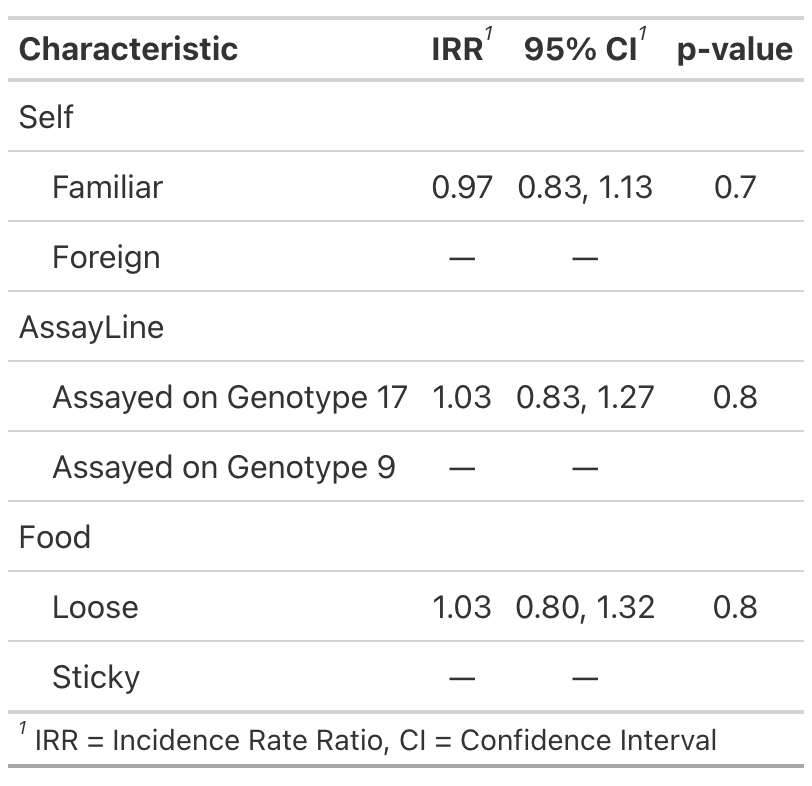 |

| **Model 3: Does food and familiar/foreign interact to affect virus phenotype in homogenous treatments?** | | |
| --- | --- | --- |
| *Data = End of evolution assays for homogeneous microcosms only* | | |
| **Composite Exploitation Rate (c)** | **Proportion Infected (i)** | **Per Infection Productivity (p)** |
| FitnessTrans ~ Self * Food + AssayLine + (1\|Treatment/Replicate/VirusLine) + (1\|Set.Up) + (1\|ObsID)  *family = nbinom1*  *package = glmmTMB* | cbind(Infected, Uninfected) ~ Self * Food + AssayLine + (1\|Treatment/Replicate/VirusLine) + (1\|Set.Up)  *family = binomial*  *package = glmer* | CountPerTrans ~ Self * Food + AssayLine + (1\|Treatment/Replicate/VirusLine) + (1\|Set.Up) + (1\|ObsID)  *family = Poisson*  *package = glmer* |
| **ANOVA**  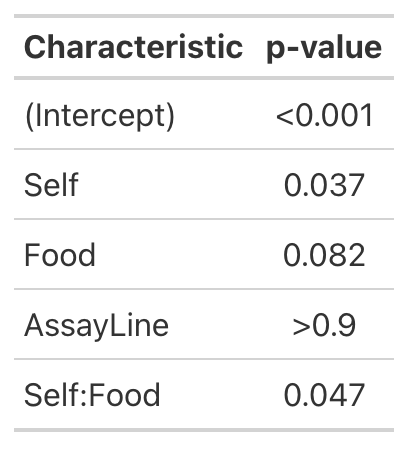 | **ANOVA**  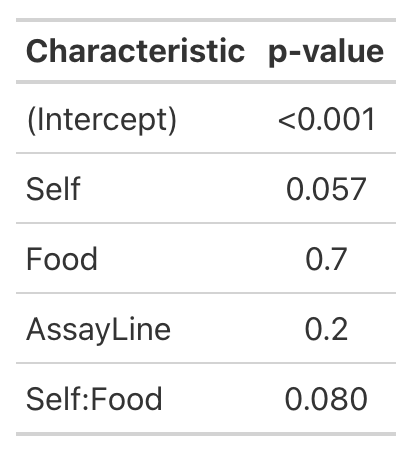 | **ANOVA**  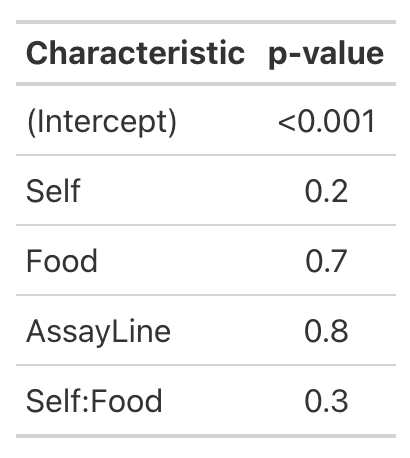 |
| **GLMM Summary**  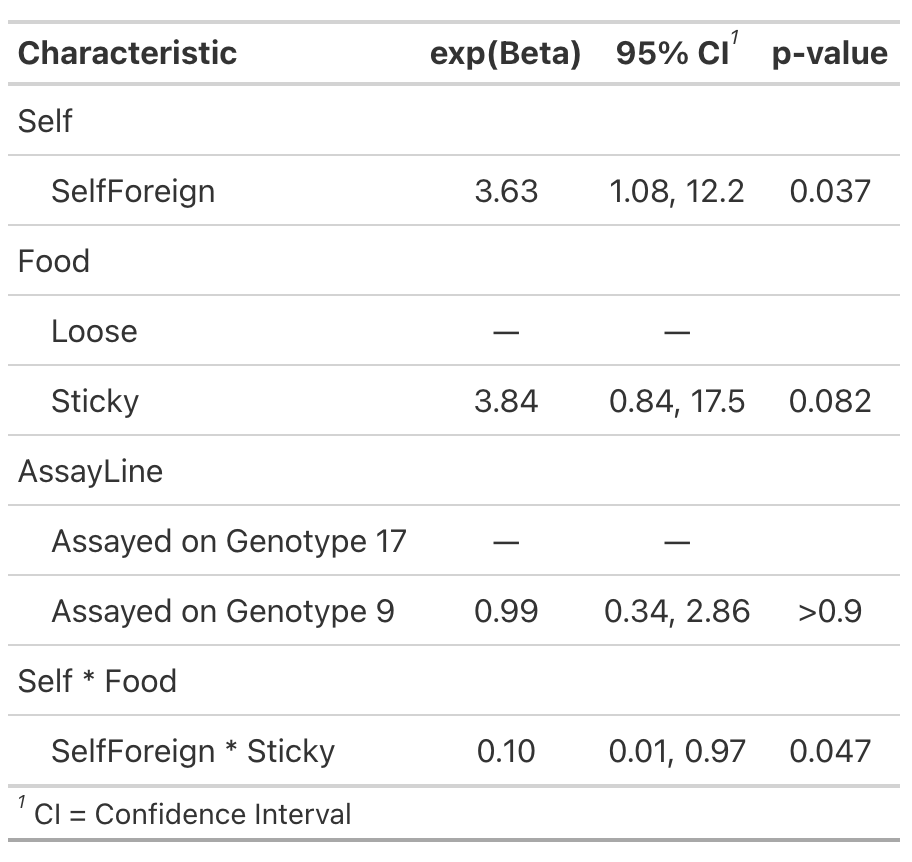 | **GLMM Summary**  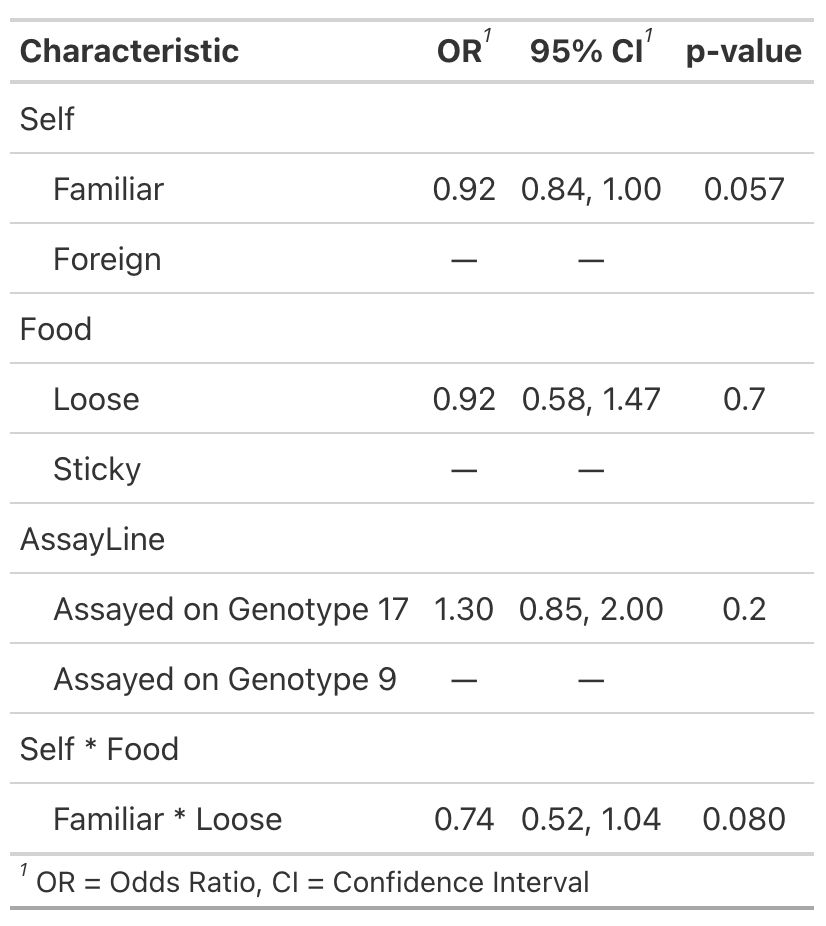 | **GLMM Summary**  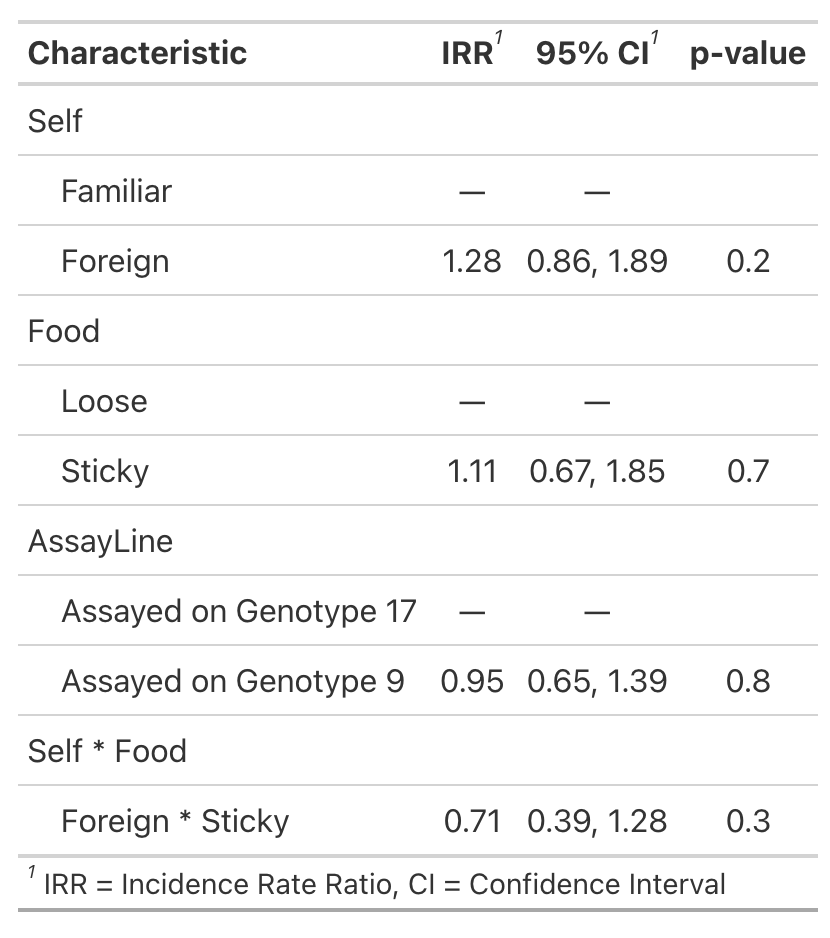 |
| **Emmeans Contrasts**  Food = Loose:  Self_pairwise estimate SE df z.ratio p.value  Familiar - Foreign -1.29 0.618 Inf -2.084 0.0372  Food = Sticky:  Self_pairwise estimate SE df. z.ratio p.value  Familiar - Foreign 1.02 0.607 Inf 1.679 0.0932  Results are averaged over the levels of: AssayLine  Results are given on the log (not the response) scale. | **Emmeans Contrasts**  Food = Loose:  Self_pairwise estimate SE df z.ratio p.value  Familiar - Foreign -0.784 0.364 Inf -2.156 0.0311  Food = Sticky:  Self_pairwise estimate SE df z.ratio p.value  Familiar - Foreign 0.442 0.359 Inf 1.233 0.2176  Results are averaged over the levels of: AssayLine  Results are given on the log odds ratio (not the response) scale. | **Emmeans Contrasts**  Food = Loose:  Self_pairwise estimate SE df z.ratio p.value  Familiar - Foreign -0.2436 0.272 Inf -0.895 0.3710  Food = Sticky:  Self_pairwise estimate SE df z.ratio p.value  Familiar - Foreign 0.0965 0.254 Inf 0.381 0.7035  Results are averaged over the levels of: AssayLine  Results are given on the log (not the response) scale. |

| **Model 4: Are heterogeneous microcosms different at a microcosm level?** | | |
| --- | --- | --- |
| *Data = End of evolution assays for all microcosms* | | |
| **Composite Exploitation Rate (c)** | **Proportion Infected (i)** | **Per Infection Productivity (p)** |
| FitnessTrans ~ HomoHetero* Food + AssayLine + (1\|Treatment/Replicate/VirusLine) + (1\|Set.Up), zi=~AssayLine  *family = nbinom2*  *package = glmmTMB* | cbind(Infected, Uninfected) ~ HomoHetero* Food + AssayLine + (1\|Treatment/Replicate/VirusLine) + (1\|Set.Up)  *family = binomial*  *package = glmer* | CountPerTrans ~ HomoHetero* Food + AssayLine + (1\|Treatment/Replicate/VirusLine) + (1\|Set.Up) + (1\|ObsID)  *family = nbinom2 (dispersion is significant)*  *package = glmmTMB* |
| **ANOVA**  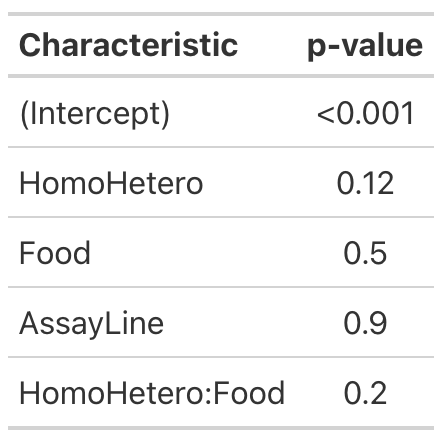 | **ANOVA**  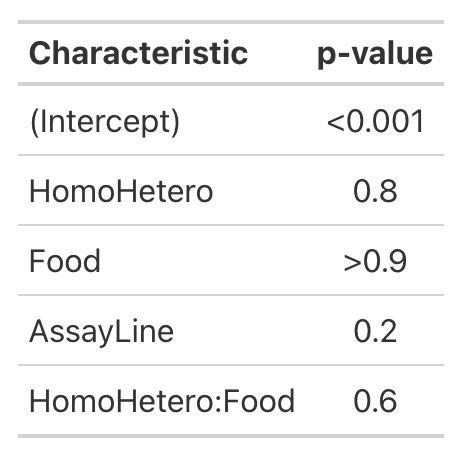 | **ANOVA**  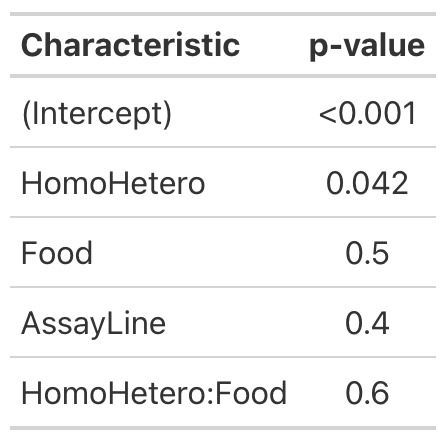 |
| **GLMM Summary**  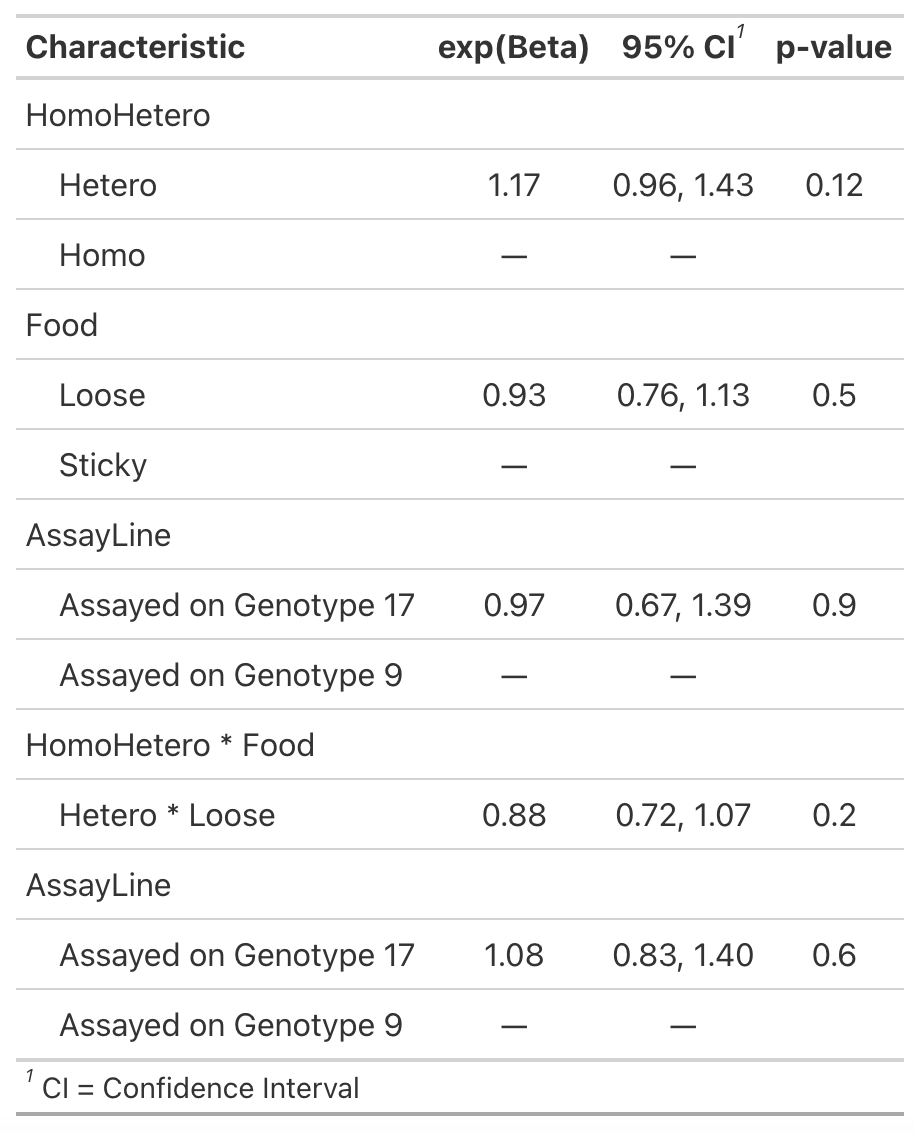 | **GLMM Summary**  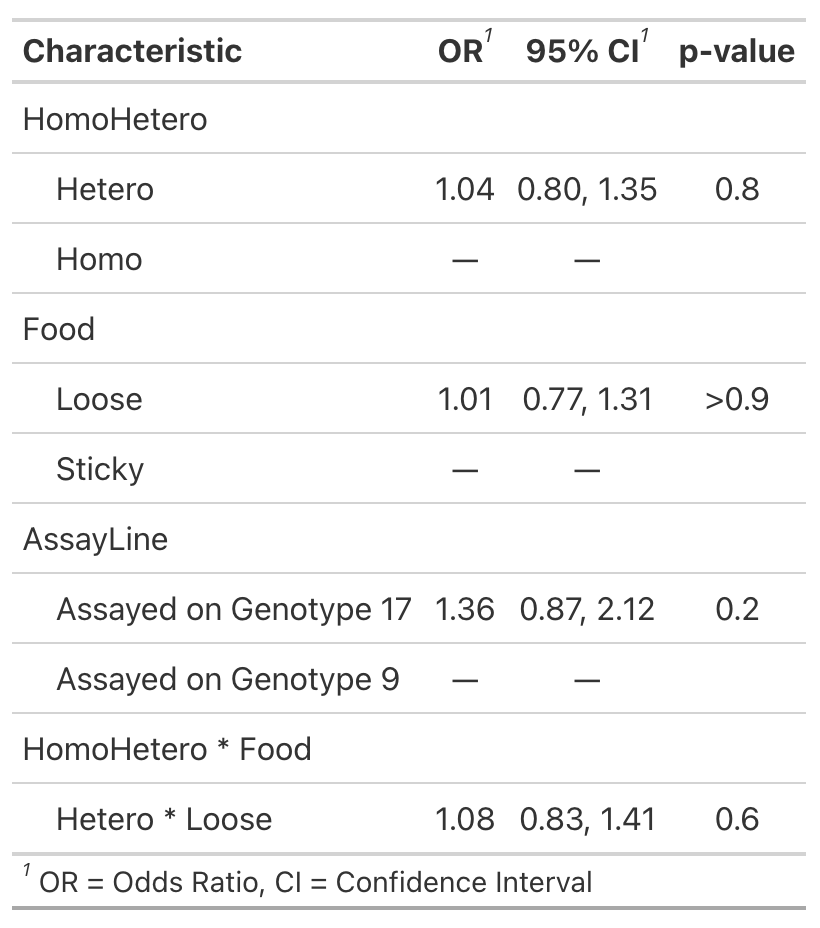 | **GLMM Summary**  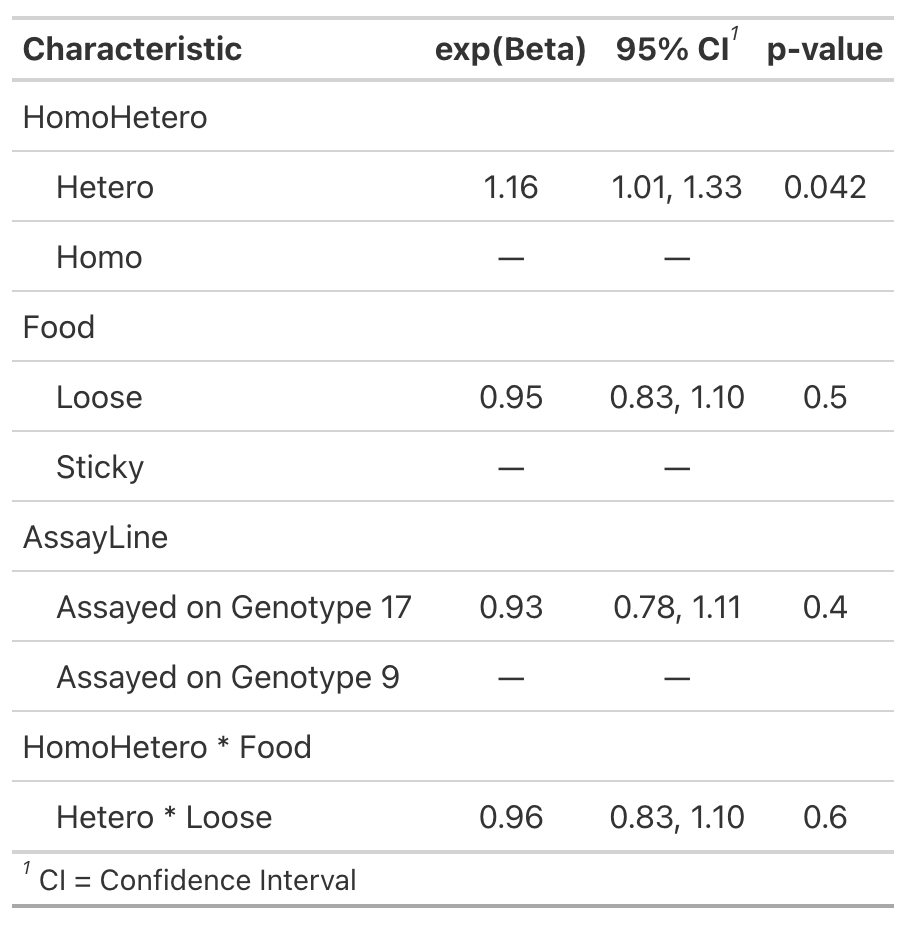 |

| **Model 5: Are the food types different at a microcosm level?** | | |
| --- | --- | --- |
| *Data = End of evolution assays for all microcosms* | | |
| **Composite Exploitation Rate (c)** | **Proportion Infected (i)** | **Per Infection Productivity (p)** |
| FitnessTrans ~ Food + AssayLine + (1\|Treatment/Replicate/VirusLine) + (1\|Set.Up) + (1\|ObsID), zi=~AssayLine  *family = nbinom2*  *package = glmmTMB* | cbind(Infected, Uninfected) ~ Food + AssayLine + (1\|Treatment/Replicate/VirusLine) + (1\|Set.Up)  *family = binomial*  *package = glmer* | CountPerTrans ~ Food + AssayLine + (1\|Treatment/Replicate/VirusLine) + (1\|Set.Up) + (1\|ObsID), zi =~AssayLine  *family = nbinom1 (dispersion is significant)*  *package = glmmTMB* |
| **ANOVA**  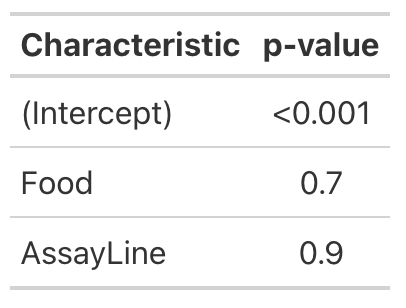 | **ANOVA**  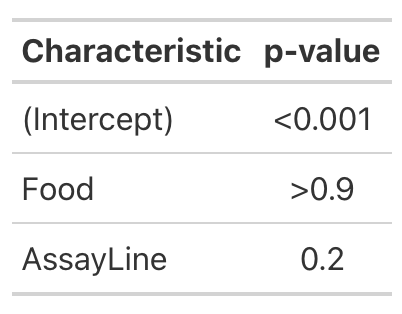 | **ANOVA**  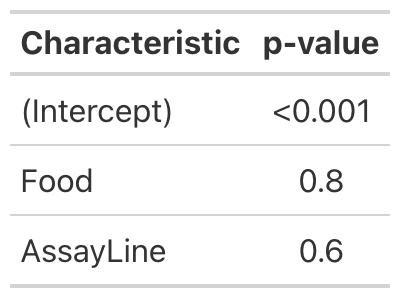 |
| **GLMM Summary**  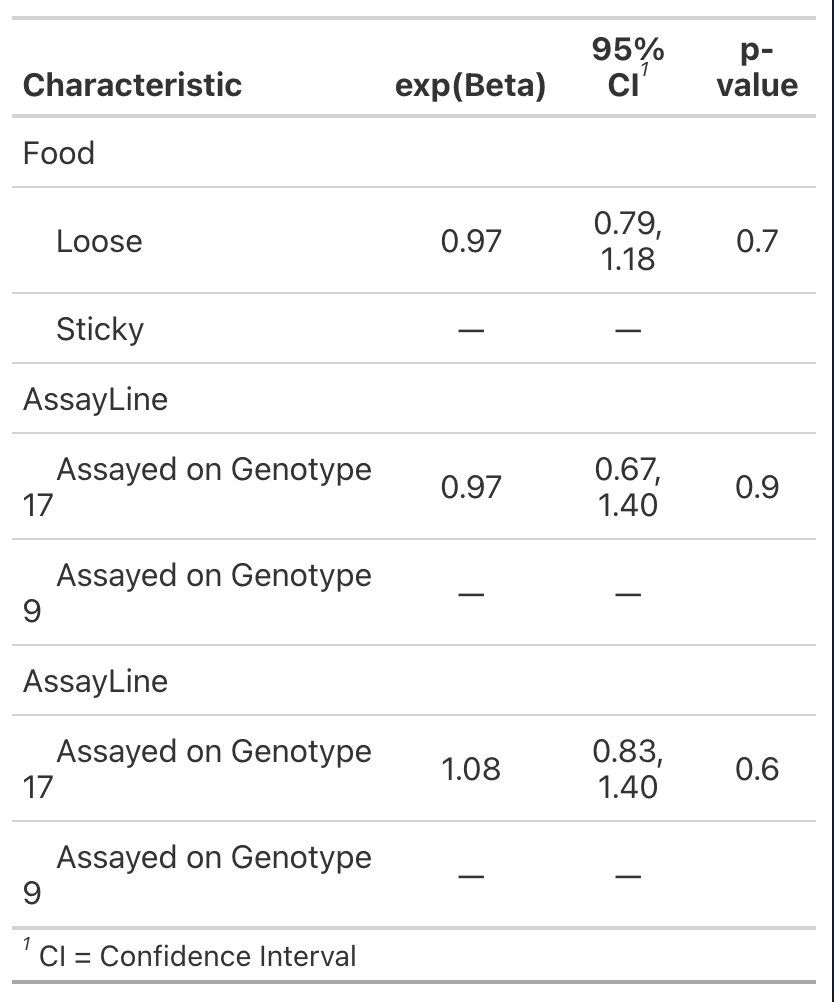  Zero Inflation | **GLMM Summary**  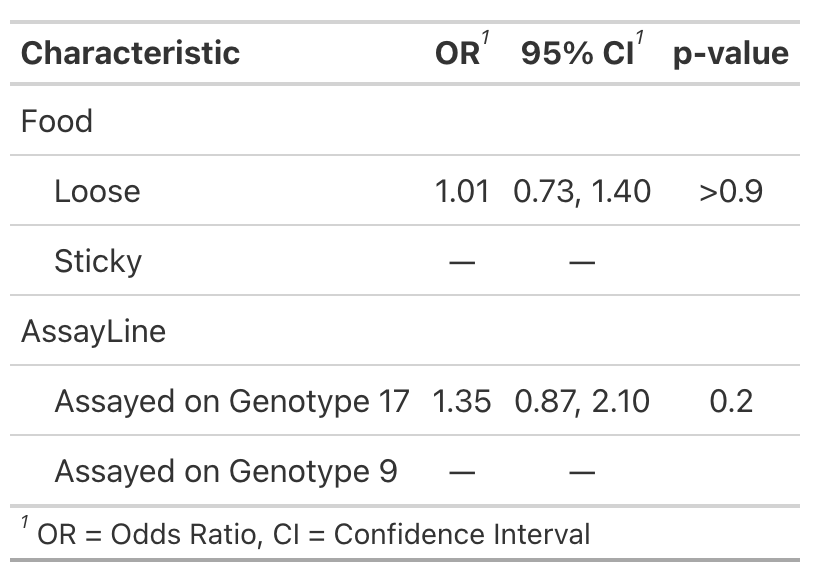 | **GLMM Summary** |

| **Model 6: Does spatial structure interact with host heterogeneity to alter virus phenotypes?** | | |
| --- | --- | --- |
| *Data = End of evolution assays for only heterogeneous microcosms* | | |
| **Composite Exploitation Rate (c)** | **Proportion Infected (i)** | **Per Infection Productivity (p)** |
| FitnessTrans ~ LocalProp17 * AssayLine * Food + Lines + (1\|Set.Up) + (1\|Position) + (1\|Treatment / Replicate / VirusLine) , zi=~AssayLine  *family = nbinom2 , package = glmmTMB* | cbind(Infected, Uninfected) ~ LocalProp17 * AssayLine * Food + Lines + (1\|Set.Up) + (1\|Position) + (1\|Treatment/Replicate/VirusLine)  *family = binomial , package = lme4* | CountPerTrans ~ Food + AssayLine + (1\|Treatment/Replicate/VirusLine) + (1\|Set.Up) + (1\|ObsID)  *family = Poisson , package = lme4* |
| **ANOVA**  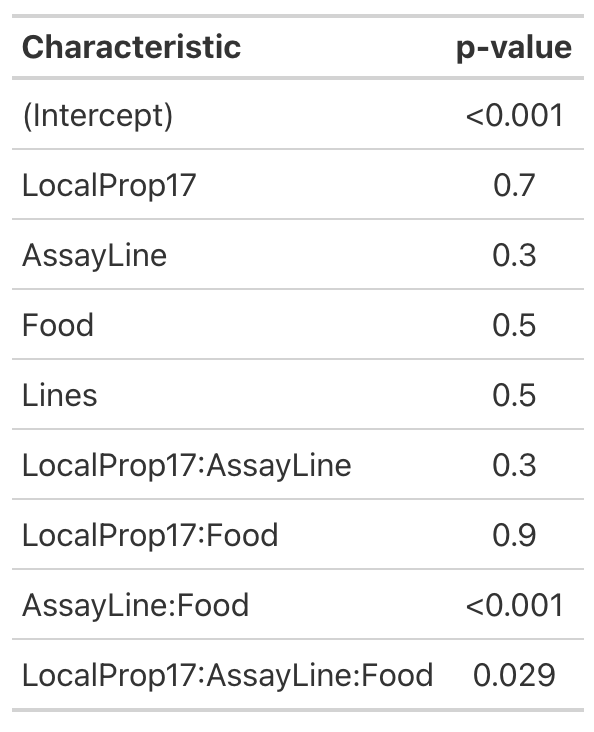 | **ANOVA**  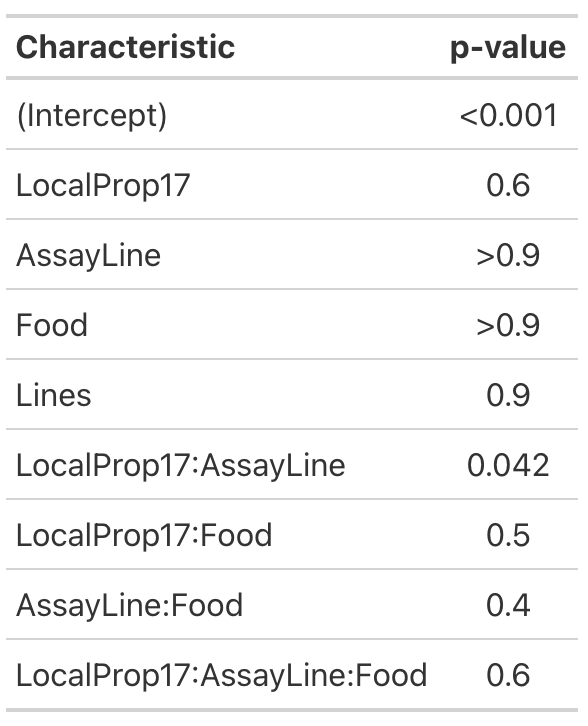 | **ANOVA**  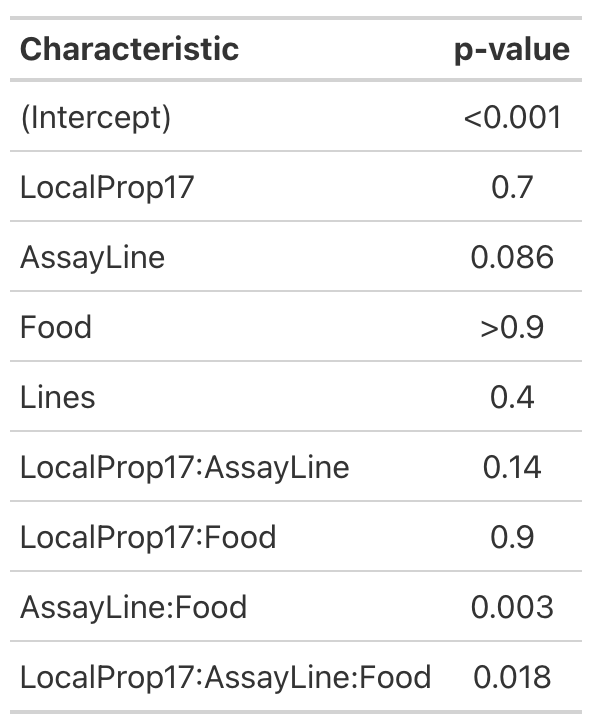 |
| **GLMM Summary**  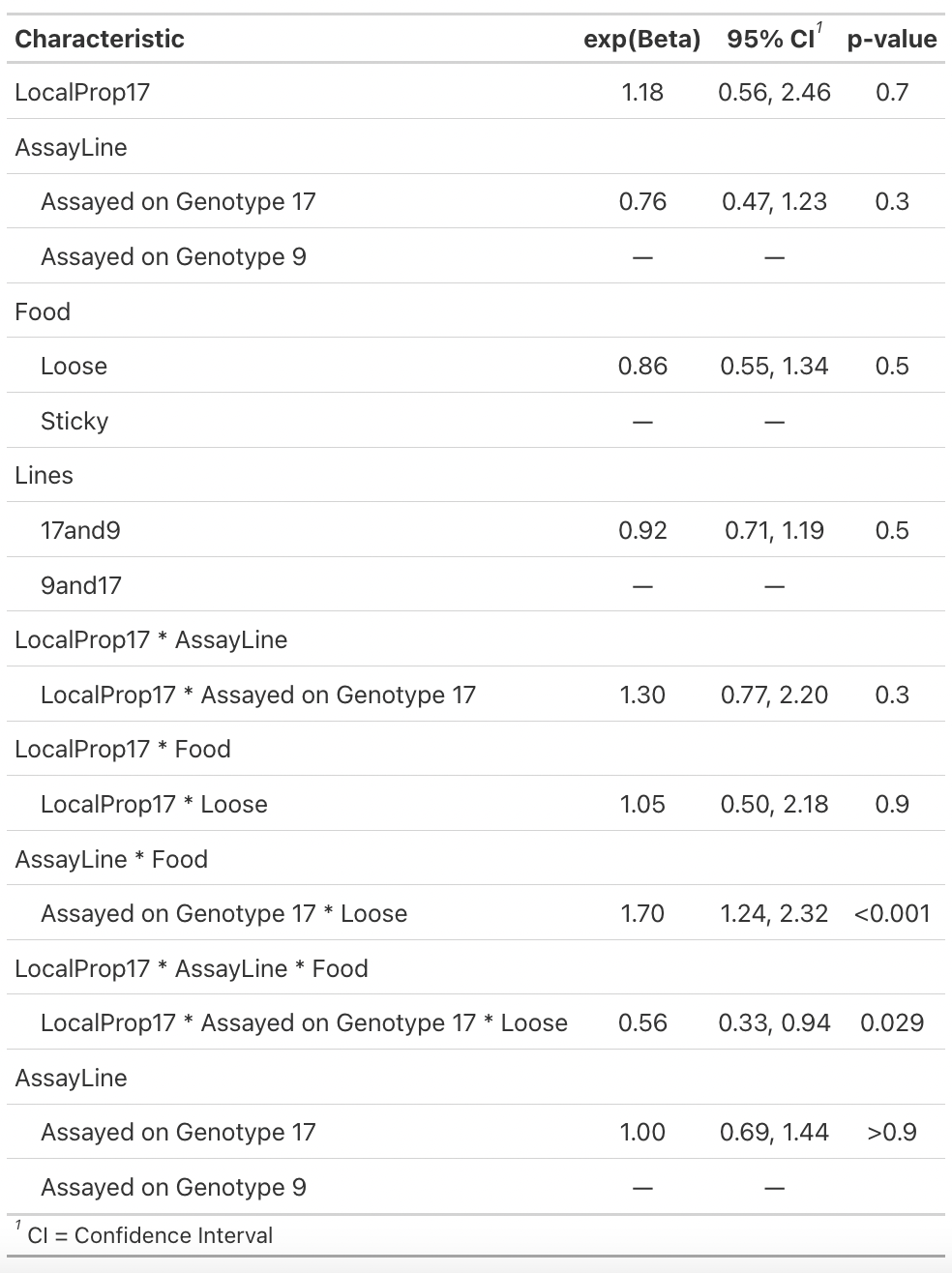  Zero Inflation | **GLMM Summary**  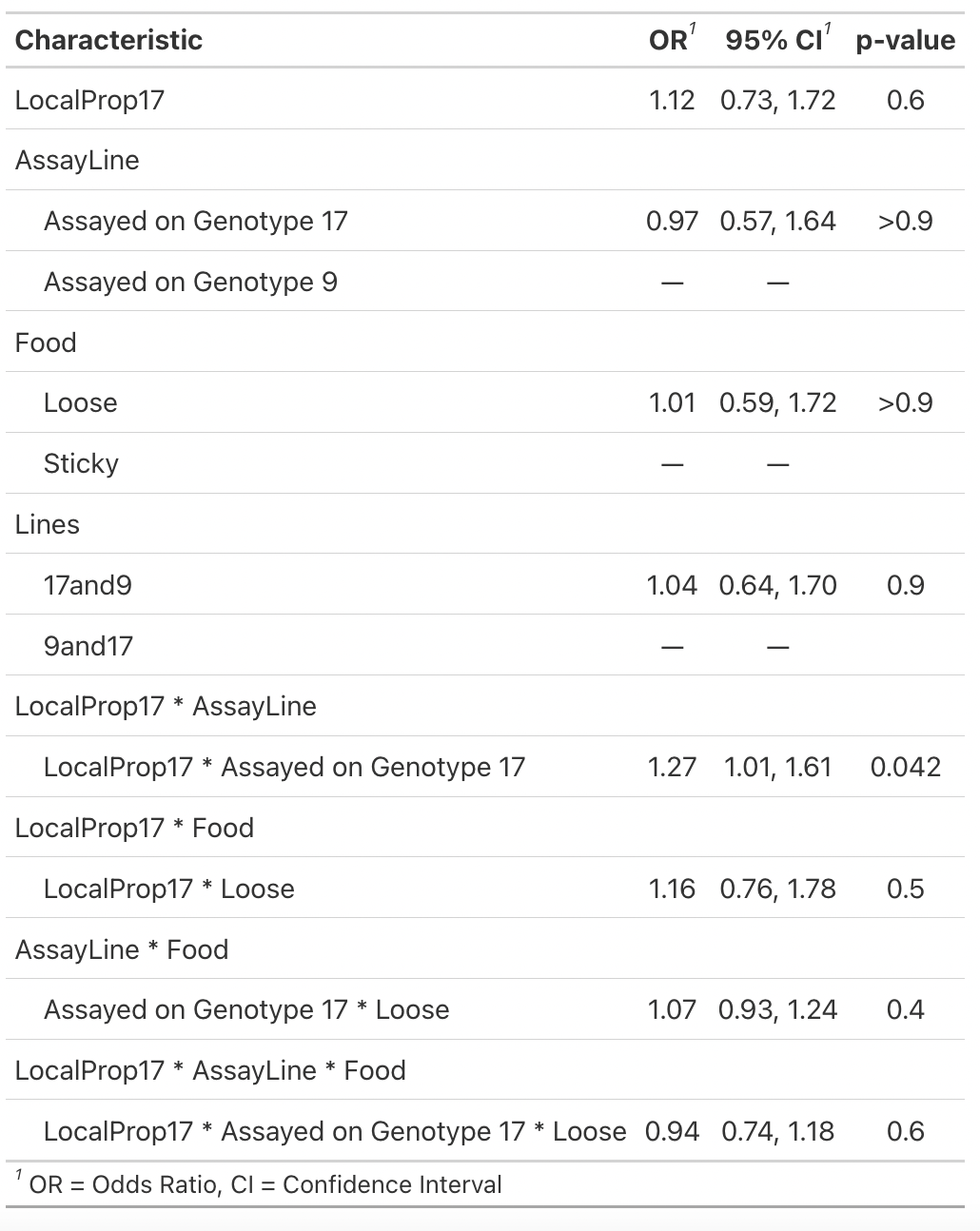 | **GLMM Summary**  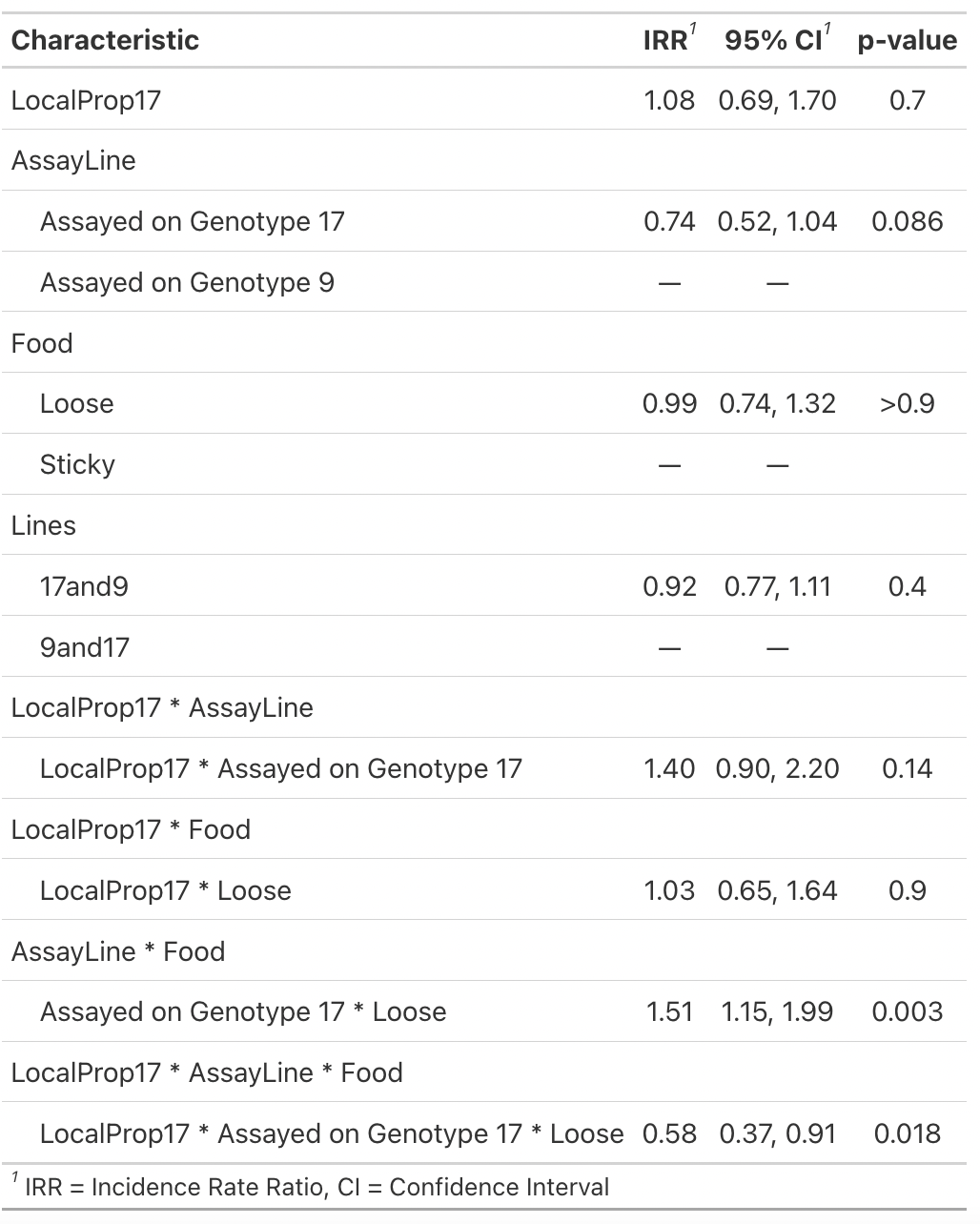 |
| **Composite Exploitation Rate EmTrends**  Model5f, ~ AssayLine*LocalProp17*Food, var = "LocalProp17", by = "Food", interaction = TRUE, adjust = 'bonferroni'  Food = Loose:  AssayLine LocalProp17 LocalProp17.trend SE df asymp.LCL asymp.UCL  Assayed on Genotype 17 0.499 -0.113 0.683 Inf -1.644 1.418  Assayed on Genotype 9 0.499 0.533 0.588 Inf -0.784 1.850  Food = Sticky:  AssayLine LocalProp17 LocalProp17.trend SE df asymp.LCL asymp.UCL  Assayed on Genotype 17 0.499 0.965 0.631 Inf -0.449 2.380  Assayed on Genotype 9 0.499 -0.735 0.697 Inf -2.298 0.829  Results are averaged over the levels of: Lines  Confidence level used: 0.95  Conf-level adjustment: bonferroni method for 2 estimates | | |
| **Composite Exploitation Rate Pair Contrasts**  pairs(Emtrends, by = "Food")  Food = Loose:  contrast  Assayed on Genotype 17 LocalProp170.5 - Assayed on Genotype 9 LocalProp170.5  estimate SE df z.ratio p.value  -0.646 0.716 Inf -0.903 0.3667  Food = Sticky:  contrast  Assayed on Genotype 17 LocalProp170.5 - Assayed on Genotype 9 LocalProp170.5  estimate SE df z.ratio p.value  1.700 0.797 Inf 2.132 0.0330  Results are averaged over the levels of: Lines | | |
